# Dissecting context-dependent cancer vulnerabilities using Perturb-seq

**DOI:** 10.64898/2026.08.24.746802

**Authors:** Samuel Maffa, Isabella A Boyle, Lie Ward, William N Colgan, Patricia Borck, Alina Simerzin, Anisah Adeagbo, Oluwatomisin Olajide, Sarah Wie, Harry Liang, Kirsty Wienand, Tsukasa Shibue, Judhajeet Ray, Brenton Paolella, Catarina D Campbell, Francisca Vazquez, Joshua M Dempster

## Abstract

**Background:** CRISPR-mediated viability assays in diverse cancer cell lines have informed cancer biology and precision medicine, but cell fitness is not the only cancer-relevant phenotype. Gene expression profiling provides insight into cellular stress, inflammation, and differential state, while still identifying activation of cell-death pathways. Perturb-seq allows scalable functional genomics screening of expression phenotypes at single-cell resolution, however existing datasets cover only a small number of work-horse cell lines.

**Results:** We produced a proof-of-concept Perturb-seq dataset targeting 100 genes in 16 diverse cancer cell lines. In the process, we established methods to address single-cell technical artifacts, identified Cas9-mediated chromosomal aberrations and assessed screen quality. Even with a limited library, we observed common signatures of deleting essential genes as well as context-specific responses based on intrinsic genomic properties of the models. For example, we inferred a previously undescribed relationship between dependence on the ER-golgi transport gene *immediate early response 3 interacting protein 1* (*IER3IP1*) and oxidative stress, demonstrating the potential of integrated Perturb-seq for hypothesis generation.

**Conclusions:** We established a framework for building a comprehensive map of post-perturbational transcriptional phenotypes using parallel Perturb-seq experiments across multiple cell lines. We demonstrated that integrated Perturb-seq experiments spanning diverse contexts enable hypotheses about gene function specific to tissue types or cancer subtypes – suggesting large-scale, genome-wide datasets would offer invaluable insight into the highly context-dependent nature of cancer biology.

## BACKGROUND

The Cancer Dependency Map (DepMap) has performed genome-wide genetic functional screening across large panels of cancer cell lines to identify genetic vulnerabilities for therapeutic exploitation [1–5]. Typically, cells engineered to express Cas protein are infected with a library of CRISPR single guide RNAs (sgRNAs) designed against a set of genes, leading to functional loss of the targeted proteins via Cas-induced frameshift mutations. CRISPR viability screens use cell recovery as the primary phenotypic readout, reporting measures of gene essentiality in a particular model system. However, this approach is blind to cell state changes due to genetic perturbations that do not produce a viability effect.

Perturb-seq experiments pair the scalability of pooled CRISPR screens with a transcriptome-wide readout of each perturbation’s effect. Similar to viability screens, cells are infected with a targeted library of sgRNAs. However, in Perturb-seq, single-cell RNA sequencing (scRNAseq) is performed to recover mRNA and sgRNA in each cell, associating transcriptomic changes with genetic perturbations. To date, Perturb-seq studies have primarily focused on a single cell line or tissue. For example, studies developing Perturb-seq and related single-cell CRISPR-based technologies [6–9] have been performed in immortalized blood cells, such as the chronic myeloid leukemia cell line K562 [10–14]. However, recent studies applying Perturb-seq to neural cells in mice have shown that the deletion of specific transcription factors affects cells in a cell-type specific manner [15–17]. In cancer biology, oncogenesis may occur in any tissue type, and even within a single lineage, diverse genetic driver events cause heterogeneous drug response [18,19]. Conversely, cancers with the same genetic driver exhibit differential drug efficacy and resistance mechanisms across lineages [20–22].

Capturing the biological diversity of cancer phenotypes requires functional assays in many contexts. Here, we generated an integrated dataset of Perturb-seq screens performed across 16 cell lines using a CRISPR Cas9 library targeting 100 genes selected to capture a variety of dependency profiles observed in DepMap. Across all cell lines, we identified then corrected for Cas9-related and other single-cell technical artifacts, and determined informative predictors of screen quality. We find that with a diverse panel of cell lines, we can distinguish universal cancer dependency phenotypes from context-specific transcriptional programs and link perturbational responses to cell-intrinsic features. In spite of our limited library, we also demonstrate the potential for novel discovery of context-dependent gene dependency mechanisms. Our work underscores the necessity of profiling multiple cell lines, highlights Perturb-seq as an information-rich complement to viability assays in functional genomics, and presents a framework for scalable screening to understand context-dependent cancer biology.

## RESULTS

### Parallel post-perturbation profiling across 16 cell lines

To understand the functional consequences of impairing selectively essential genes in different cancer models, we performed Perturb-seq experiments in 16 Cas9-engineered cancer cell lines with a lentiviral CRISPR guide library targeting 100 genes (Figure 1a). We chose gene targets from four classes of fitness effect distributions using the DepMap CRISPR data: positive controls (commonly essential genes), strongly selective (dependent in only a minority of cell lines), high variance across cell lines, and negative controls (unexpressed olfactory receptors) (Figure 1b, Supplementary Table 1). Selective and high variance dependencies are interesting as the range of responses seen in DepMap might suggest a potential therapeutic window [23]. We included both highly predictable and poorly explained dependencies (Supplementary Figure 1a). We selected these targets with the aim of gaining additional insight into their dependency patterns through post-perturbational expression profiling.

**Figure 1.**
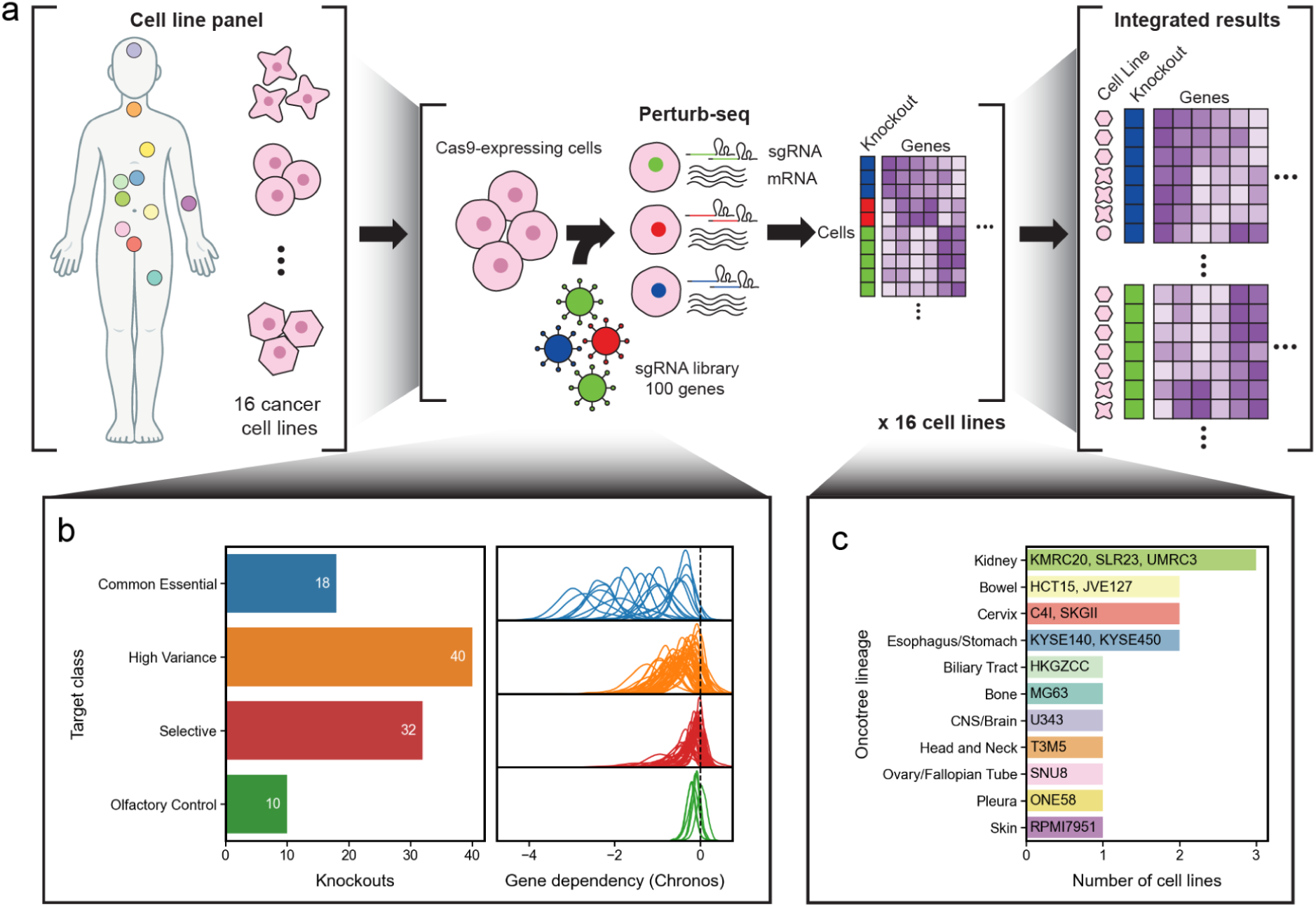
Overview of experimental design. **a.** Perturb-seq screens using Cas9 knockout were performed independently in 16 human cancer cell lines, representing a variety of tissues. Single-cell expression data across all screens were subsequently integrated across all models. **b.** Target genes for the CRISPR library were selected from different dependency classes across all DepMap cell lines, with olfactory receptor genes serving as non-essential negative controls. **c.** The panel of screened cell lines, covering 11 distinct lineages of origin.

In tandem with designing the target library, we selected a panel of 16 cell line models representing 11 lineages (Figure 1c, Supplementary Table 2), ensuring that the majority of selective and high variance dependencies exhibited a strong cell viability effect in at least one included cell line. Most cell lines are wild-type for target genes in our library, although some exceptions include amplification of *MYC* and *AP2M1* mutations (Supplementary Figure 1b). We constructed our target library and cell line panel such that our knockouts showed selective dependency in different subsets of cancer models (Supplementary Figure 1c). We identified single guide knockouts using SCEPTRE low-MOI (see Methods) [24].

### Identification of CRISPR-induced artifacts

We first wanted to evaluate the quality of the dataset and identify any potential biases or artifacts. For example, Cas9-mediated double-stranded breaks are known to cause unwanted bystander effects, including chromosomal arm truncations [25–27] and corresponding partial arm gain [28]. In principle, single-cell data can identify these chromosomal aberrations at a per-cell level. We identified Cas9-induced truncations or gains using the average expression in all genes distal to the target gene on the same chromosome arm. Some cells with negative control (olfactory receptor) knockouts exhibited position-dependent loss of expression consistent with arm truncation, while others showed increased expression (Figure 2a). Across all control knockouts in all cell lines, we found decreased expression at the targeted arm (median z-scored expression = −0.022), which was ameliorated by removal of cells with aberrant position-dependent arm expression (median z-scored expression = −0.0003, Figure 2b). Chromosome arm breaks can be used to identify screen-specific off-target editing. For example, *OR11H1* vectors cause truncation not only at *OR11H1* but also on chromosome arm 14q, which contains an olfactory gene cluster near its centromere [29] (Figure 2b).

**Figure 2.**
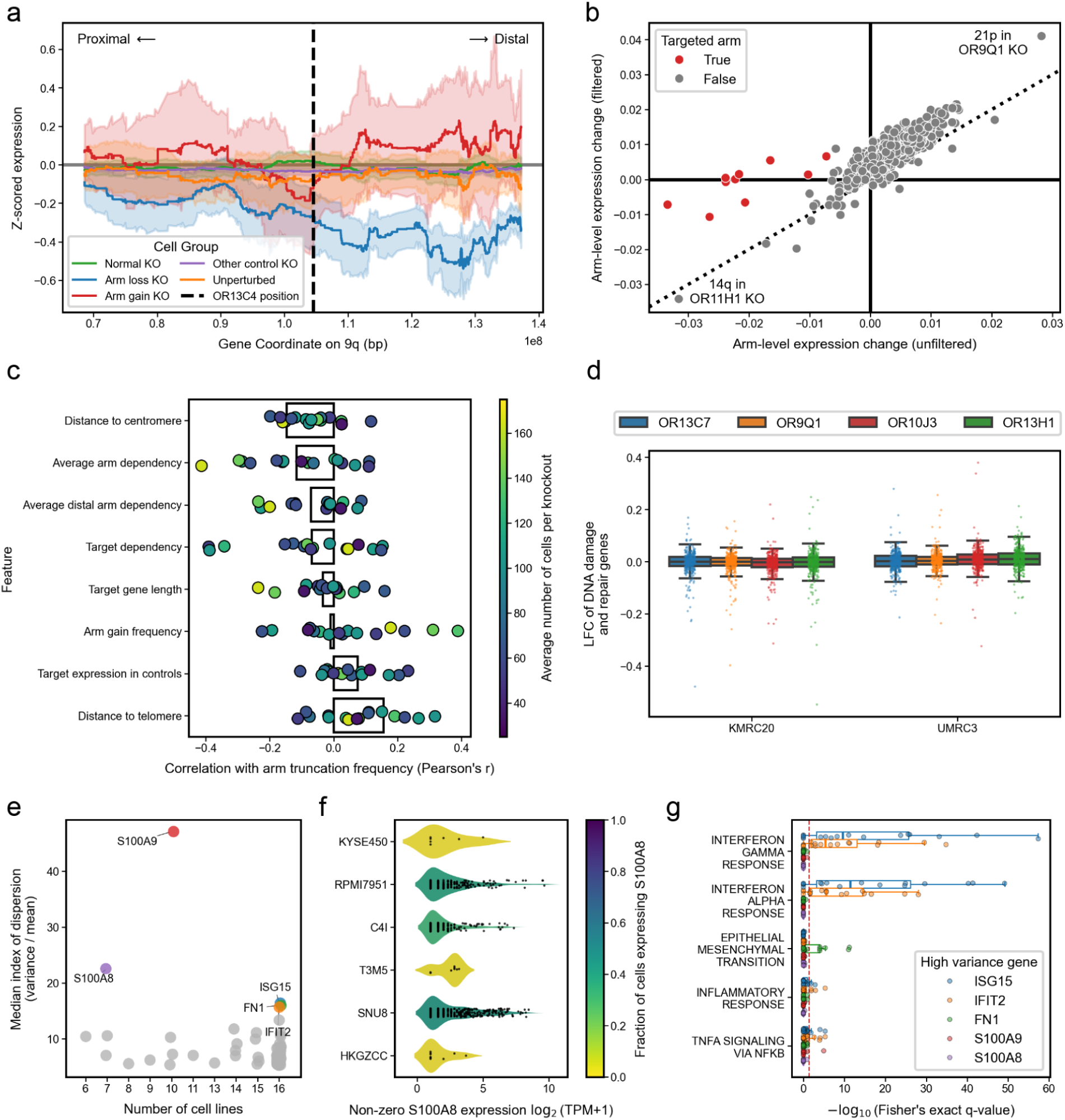
Artifacts of CRISPR knockout and examination of cell heterogeneity. **a.** Average z-scored expression along the targeted chromosome arm in cells with a negative control knockout, *OR13C4*, classified as normal ploidy, arm truncation, or arm gain. The averages from other knockout and nontargeting controls are shown for reference. Error bands represent 95% confidence intervals. **b.** Average change in expression over chromosome arms between cells receiving a control knockout compared to all other control knockouts, before and after filtering cells for putative arm loss. **c.** Correlations between gene-specific properties and frequency of arm truncation. Bars represent relationships calculated over all combinations of cell lines and knockouts; points represent relationships calculated over individual cell lines. **d.** Log-fold change in expression of DNA damage and repair genes in olfactory control knockouts compared to non-cutting controls. **e.** Consistently high variance genes ranked below the 97th percentile in average expression **f.** Violin plot of non-zero gene expression of *S100A8* in individual cells from expressing cell lines. ONE58 is not shown due to having a single cell with non-zero expression. Colored by fraction of cells with any S100A8 expression. **g.** Enrichment of selected Hallmark gene sets among the top correlates of the five most variably expressed genes with average expression below the 97th percentile in negative control cells.

We noted that the frequency of arm loss varied by both gene and cell line. The consistency of arm truncation frequency between constructs targeting the same gene (Pearson’s *r* = 0.46, *p* = 8.96 x 10^−64^, *n* = 1,194) suggests that arm truncation rates were related to properties of the genetic locus, rather than the guide sequence (Supplementary Figure 2a). We assessed several gene-specific properties, including distance to the centromere, expression of the target gene, and essentiality of the targeted arm, but found no clear feature that explained the frequency of arm loss over different genes (Figure 2c). When evaluating predictors in cell lines individually, 4 cell lines exhibited a modest relationship (Pearson’s *r* between −0.25 and −0.41) where chromosome arms with stronger gene-wise fitness effects on average paradoxically show a higher frequency of arm loss, but this trend is weaker when calculated over all cell lines together (Pearson’s *r* = −0.12, *n* = 1,531) or the remaining 12 models individually (Pearson’s *r* between −0.2 and 0.2, Figure 2c, Supplementary Figure 2b). It’s possible that the causal arrow points the other way: gene knockouts that frequently cause arm truncation events will appear to lower cell viability, independent of the specific targeted gene. A previous study by Lazar et al. reported that the presence of functional *TP53* affects the frequency of arm truncation [25], but we find no such relationship across our cell lines, possibly because our dataset is limited to only 16 models (median frequency in *TP53* mutant = 14%, *TP53* wild-type = 12.5%; Mann-Whitney *p* = 0.31, Supplementary Figure 2c).

In general, we observed fewer putative arm gain events than arm loss events (median percentage of cells per perturbation with arm gain = 3.7%; arm loss = 13.1%). Across all perturbations, the frequency of arm gain was weakly correlated the essentiality of the target gene (Pearson’s *r* = −0.21, *p* = 2.35 x 10^−17^, *n* = 1,531), but moderate correlation (Pearson’s *r* > 0.3) between arm gain and arm loss was only observed in JVE127 and ONE58 (Supplementary Figure 2d). Among cells receiving control knockouts, we found that *SGO1-AS1* expression was lower in cells with arm gain, but more highly expressed in cells with arm loss (p=1.68 x 10^−5^, Supplementary Figure 2e). *SGO1-AS1* is a repressive antisense transcript to a gene required for chromosome segregation [30]. However, a larger library of targets would be necessary to determine the mechanisms behind arm truncation frequency. Our use of dual-sgRNA vectors may have increased arm aberration events, as each vector has two guides causing double stranded breaks in close proximity.

Even when Cas9-induced double-strand breaks do not cause large genomic losses, they might still cause genotoxic stress that distorts the transcriptional response of cells to gene knockout. We evaluated this possibility by rescreening UMRC3 and KMRC20 with a limited library of 4 olfactory receptor control knockouts (4 guides per knockout) and 4 non-targeting guides but great depth (63,000/42,000 mean UMIs per cell, 4 x 10^7^/6 x 10^7^ mean UMIs per knockout respectively). Using SCEPTRE low-MOI after filtering out cells with detected arm artifacts, we tested for differentially expressed genes between cutting and non-cutting controls. All genes called as significant (FDR < 0.1) had either small effect size or low expression level, indicating a negligible difference in transcriptional state following the Cas9 induced-double strand break (Supplementary Figure 2f). Critically, we did not observe any signs of genotoxic stress as measured by no change in expression of genes in Reactome DNA damage/repair pathways in the cutting similar to non-cutting controls (Figure 2d).

We next excluded cells based on a low number of unique molecular identifiers (UMIs), genes detected, aberrant proportion of mitochondrial reads, inability to assign single guide infections, and arm truncation. Overall, the recovery rates of high-quality singly-infected cells varied greatly by cell line (22-54%), where sequenced cells were commonly excluded due to having ambiguous CRISPR infections based on mixed sgRNA counts with low abundance (17-66%), no detectable infection (.001-30%), or poor gene expression recovery (2-10%). Overall chromosomal arm aberrations affected 10-22% (median = 17%) of cells otherwise passing quality control (Supplementary Figure 2g-h). We also compared cell line pseudobulk expressions with their DepMap bulk RNASeq profiles and confirmed that they matched (see Methods, Supplementary Figure 3a).

### Examination of cell heterogeneity

The single cell resolution in Perturb-seq provides an opportunity to observe heterogeneous cell programs within a screen. Given that cancer is highly plastic[31], we examined whether transcriptional variation was observable in negative control cell populations. We identified 321 high variance genes with median index of dispersion above the 99th quantile and present in more than 5 cell lines. The high variance genes were generally highly-expressed (*n* = 241 above the 97th percentile expression), including mitochondrial genes (*n* = 10), ribosomal proteins (*n* = 77), and housekeeping genes (*n* = 121) [32,33]. However, 80 of these highly variable genes ranked below the 97th percentile in average expression (Figure 2e). For example, *S100A8* displayed remarkably high variance despite only being detected in seven cell lines and in at most 44% of cells in any cell line (Figure 2f).

We posited that these high variance genes may serve as markers of rare cell programs, independent of perturbation. To gain a more comprehensive view of these programs, we identified other genes that are highly correlated with each of the top five outlier high-variance genes with average expression below the 97th percentile (*S100A9*, *S100A8*, *ISG15*, *FN1*, *IFIT2*) and tested for gene set enrichment among their top correlates. Notably, the genes correlated with *ISG15* and *FN1* were enriched for interferon response genes and epithelial-mesenchymal transition genes respectively (*q* < 5 x 10^−3^, Figure 2g). Considering control cells represent a theoretically homogenous monoculture in these screens, the heterogeneity of these functional clusters lends support to the picture of pro-inflammatory and epithelial/mesenchymal states as intrinsically variable and highly dynamic. Interestingly, top correlates of *S100A8* and *S100A9* display no apparent enrichment for related Hallmark gene sets despite their reported roles in modulating the inflammatory response (Figure 2g) [34], nor are *S100A8* or *S100A9* expression levels strongly correlated with *ISG15* and *FN1*.

### Quality control of perturbational profiles

Next, we evaluated the viability effect of gene knockouts using cell recovery. Although our screens are 7 days rather than the 21 days of most DepMap viability screens, we expected more negative gene effects in DepMap to correspond to lower recovery of cells in Perturb-seq. We found a strong correlation between cell depletion and DepMap gene effect scores irrespective of cell line identity (Figure 3a, Pearson’s *r* = 0.64, *p* = 3.1 x 10^−177^*, n* = 1,525) and within each individual cell line (Supplementary Figure 3b, Pearson’s *r* between 0.5 and 0.78; median = 0.66), indicating that the Perturb-seq experiments corroborate prior viability screening data for these models.

**Figure 3.**
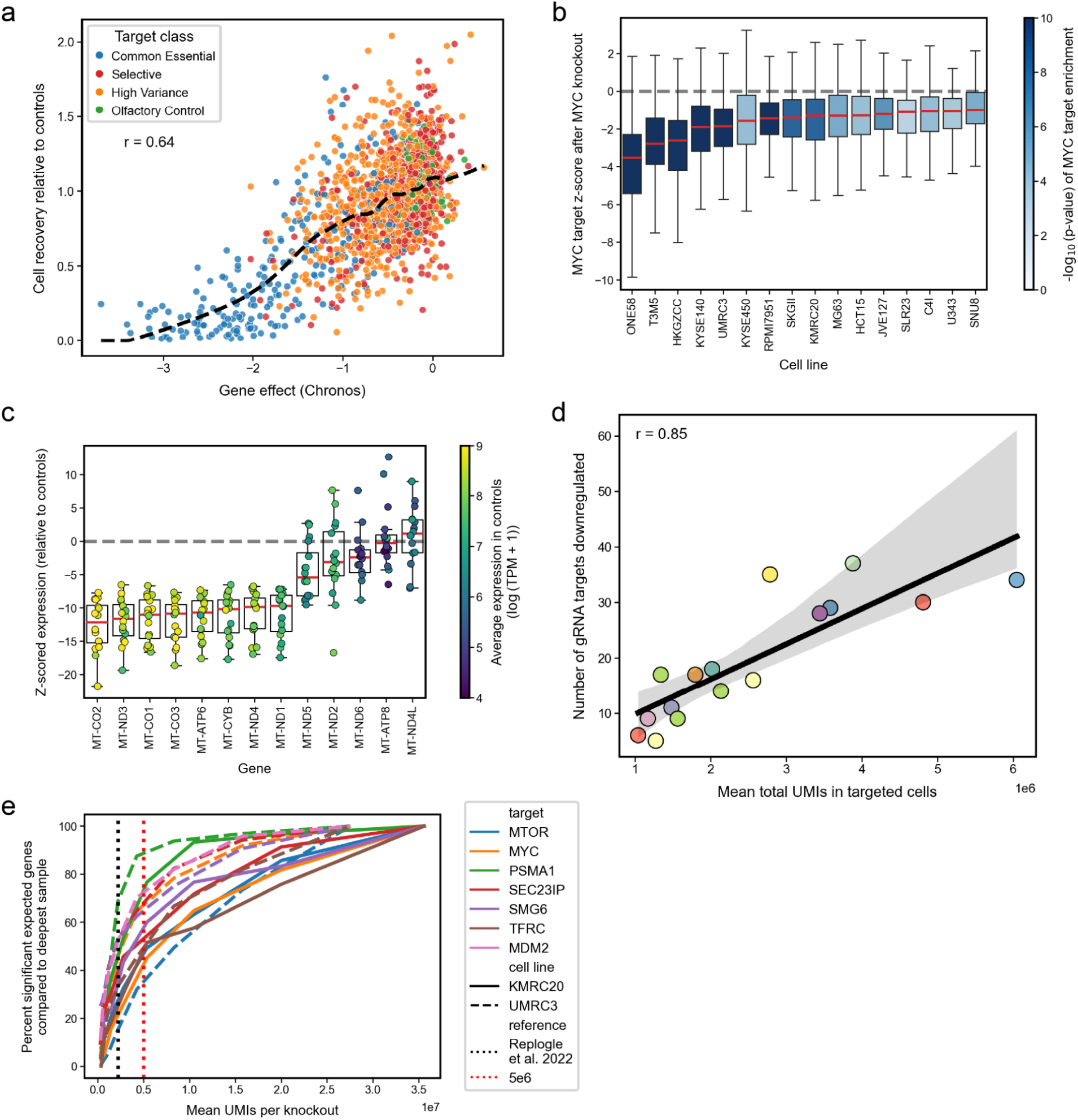
Quality control of Perturb-seq screens. **a.** Cell recovery per knockout in our screens vs CRISPR gene viability effect as reported in DepMap 25Q2. Dashed line represents the LOWESS regression trend. **b.** Z-scored expression change of genes in Hallmark MYC targets per cell line following MYC KO, colored by gene set enrichment of the combined set of *MYC* targets. **c.** Z-scored expression change of mitochondrial genes following *MTPAP* knockout across all cell lines, colored by average expression level in control cells. **d.** The number of gene targets detected as significantly downregulated per screen vs average total UMIs per perturbation. Black line represents trend with 95% confidence intervals. **e.** Recovery of expected signatures in downsamples of deep rescreens as a percent of those detected in the deepest downsample.

While cell dropout can be studied in ordinary viability screens, Perturb-seq allows researchers to study differential gene expression after perturbation at scale. The volume of reliable gene expression changes detected is one measure of the signal provided by a Perturb-seq experiment. We performed differential expression analysis with SCEPTRE, comparing cells with test gene knockouts to cells with negative control olfactory knockouts in each cell line. We expected that most Cas9 knockouts would reduce the targeted gene’s own transcripts through nonsense mediated decay, and found most of the target genes showed significant downregulation (61 / 64 with *q* < 0.05). The three exceptions where knockout consistently led to upregulation of the targeted gene were *YTHDF2*, *MYC*, and *MDM2*; this possibly indicated a compensatory response to nonfunctional protein products (Supplementary Figure 3c). Across all screens, we observed the expected downregulation of MYC targets following *MYC* knockout (Figure 3b), a further indication of data fidelity. *MTPAP* provided an additional positive control for transcriptional phenotype: knockout of *MTPAP* leads to loss of polyadenylation of mitochondrial genes, causing a subset to show severe depletion in poly-A capture methods such as used here (Figure 3c).

To assess statistical power more systematically, we used the number of significantly downregulated targets strongly as a proxy for captured signal (Supplementary Table 3). This metric was highly correlated with the total number of UMIs recovered across all cells (Pearson’s *r* = 0.85*, p* = 3.73 x 10^−5^, *n* = 16; Figure 3d). In contrast, commonly reported metrics of quality in single cell experiments, such as the average number of UMIs per cell (Pearson’s *r* = 0.45, *p* = 0.08), the number of cells per condition (Pearson’s *r* = 0.34, *p* = 0.2), and the average number of unique genes detected per cell (Pearson’s *r* = 0.54, *p* = 0.3) were less strongly correlated with target downregulation (Supplementary Figure 3d-f). Based on these results, we concluded the strongest predictor of Perturb-seq statistical power was the total number of UMIs recovered in the experiment. We downsampled the two deeply sequenced rescreens of UMRC3 and KMRC20 to assess how recoverable the signal for seven well characterized knockouts varies with total UMIs captured. The answer varied by target, with an elbows around 5 million UMIs/perturbation (Figure 3e) – a reasonable target for future screens, at nearly double the depth used in the gold standard Replogle et al. dataset [10]. However, saturating the recovery of expected differential expressions may require more than 10 million UMIs/perturbation.

### A global survey of transcriptional responses in cancer dependencies

Post-perturbational transcriptomics can resolve viability loss into distinct stress phenotypes. We observed stronger deviations from a cell line’s basal transcription state as the strength of the dependency increased (Pearson’s *r* = −0.67, *p* = 3.16 x 10^−195^, *n* = 1,523; Figure 4a). In dependent cell lines, selective and high variance targets induced expression changes commensurate with pan-essential targets.

**Figure 4.**
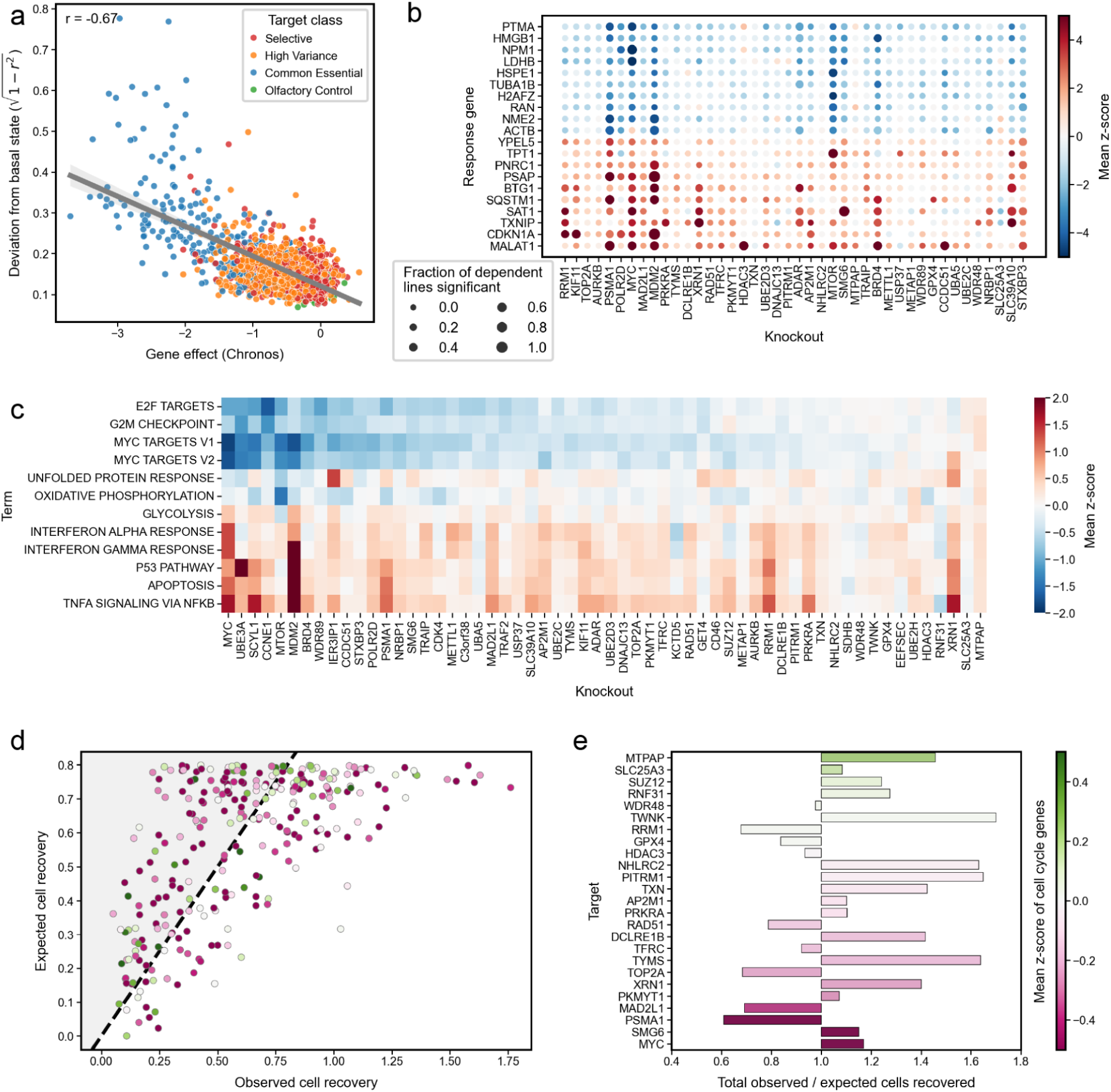
Global analysis of dependent perturbations (gene effect < −1) across all cell lines. **a.** Deviation from basal expression state vs DepMap CRISPR gene effect for each knockout. The grey line represents the trend with 95% confidence intervals **b.** Dotplot of the most commonly differentially expressed genes across all dependent (DepMap gene effect < −1) perturbations. **c.** Mean expression change per Hallmark pathway across dependent cell lines per knockout. **d.** Expected cell recovery relative to control cells by regression on DepMap gene effect scores compared to observed cell recovery for dependent perturbations (N = 313). Dashed line represents y=x. **e.** Ratios of cell recovery compared to expected recovery per knockout. Colors in **d** and **e** represent the mean transcriptional change of the Hallmark G2M_CHECKPOINT and E2F_TARGET pathway genes.

To investigate common markers of cell stress or early markers of cell death, we identified the most frequent differentially expressed genes across knockouts in dependent genes (Figure 4b). In general, we observed more consistency in the transcripts that were downregulated upon knockout of a dependent gene (10th percentile correlation = 0.49, Supplementary Figure 4a), with the top 50 most strongly downregulated genes enriched for MYC targets, pathways related to viral infection, and translation (*q* < 0.05 x 10^−9^, Supplementary Figure 4b). In contrast, there was more variability in the commonly upregulated transcripts (10th percentile correlation = 0.13, Supplementary Figure 4a), which were enriched for apoptotic pathways, hypoxia, and FOXO-mediated transcription (*q* < 0.05 x 10^−4^, Supplementary Figure 4b).

We additionally measured broader signatures of cell stress response by computing average expression changes at the pathway level, using the highly curated Hallmark collection of gene sets [33]. We found a gradient of downregulation in Hallmark pathways associated with cell cycle progression across dependent knockouts (*n* = 58), such as G2M_CHECKPOINT and E2F_TARGETS [33], which are correlated with MYC target gene sets (Pearson’s *r* > 0.62, Supplementary Figure 4c). Additionally, we observed modest upregulation of apoptotic pathways across most dependent knockouts (median z-score > 0.25, Figure 4c). Both observations are consistent with findings at the individual gene level and suggest that cell cycle arrest and apoptosis are the most common convergent phenotypes associated with a loss of fitness. The interferon pathways and the apoptotic pathways showed correlated responses across dependent knockouts (mean Pearson’s *r* = 0.57). However, gene sets such as the unfolded protein response (UPR) and oxidative phosphorylation exhibited knockout-specific responses uncorrelated with apoptotic signatures (Pearson’s *r* = 0.012, −0.091, respectively) but modestly related to cell proliferation (Pearson’s *r* = 0.21, 0.35, respectively, Figure 4c, Supplementary Figure 4c).

Although most dependent knockouts downregulate cell cycle markers, a few counterintuitively upregulate cell cycle regulatory genes in the Hallmark E2F_TARGETS and G2M_CHECKPOINT gene sets. Targets in this category include *PITRM1*, *NHLRC2*, *MTPAP*, *PRKRA*, *RNF31*, *SUZ12*, and *SLC25A3*. We also recovered more cells for these knockouts than expected given the strength of the dependency reported in DepMap (Figure 4d). Given that strong viability effects have been observed in genome-wide screens in the same cell lines after 21 days, we hypothesize that these knockouts are slower dependencies where the viability loss phenotype emerges sometime after the 6-day timepoint used in these Perturb-seq screens (Figure 4e). We looked for hints of future loss of viability in the differentially expressed genes following these knockouts using Gene Ontology’s Biological Process gene sets [35,36]. The strongest signals included oxidative phosphorylation in *MTPAP* knockout (*q* = 9.74 x 10^−19^), immune response in *RNF31* knockout (*q* = 6.81 x 10^−8^), and apoptosis in *SUZ12* knockout (*q* = 0.020) (Supplementary Figure 4d). The remaining knockouts did not induce a significant change in expression, with no gene exhibiting differential expression except for *SLC25A3* in its own knockout.

### Identification of common and context-dependent transcriptional responses

Perturb-seq can help label gene function using guilt by association, i.e. finding gene knockouts with similar transcriptional responses to a query knockout. With diverse cancer cell lines, we can extend guilt by association across cellular contexts. We defined a network of gene targets using pairwise correlations between Z-score change in expression of top differentially expressed genes (set of top 25 significant response genes at FDR < .05 per perturbation in perturbations with at least 5 significant responses, n=2819) across our entire integrated dataset spanning 16 cell lines and 90 perturbed genes, excluding perturbations for which we recovered fewer than five cells (*n* = 1,394, Supplementary Figure 5; Figure 5a). Filtering edges to highly correlated knockout pairs (Pearson’s *r* > 0.5) across any four pairs of cell lines, we observed two clusters which exclusively showed similarity in disjoint sets of cell lines, while the remaining network included edges with high similarity between knockouts in the same cell lines (Figure 5b). Some knockouts induced similar phenotypes in almost all cell lines. For example, the context-independent cluster, *XRN1* and *SMG6* showed the highest degree of similarity across all pairs of cell lines (0.18 < Pearson’s *r* < 0.70, median = 0.43), with *ADAR* knockout inducing a similar response (median Pearson’s *r* = 0.23, 0.22 to *XRN1* and *SMG6*, respectively, Supplementary Figure 6a). Both *XRN1* and *SMG6* perturbations induced an extreme upregulation of small nucleolar RNAs (13 / 32, 14 / 32 with z-score > 5, respectively), consistent with their roles as nucleases in the nonsense-mediated decay pathway (Figure 5c, Supplementary Figure 6b) [37].

**Figure 5.**
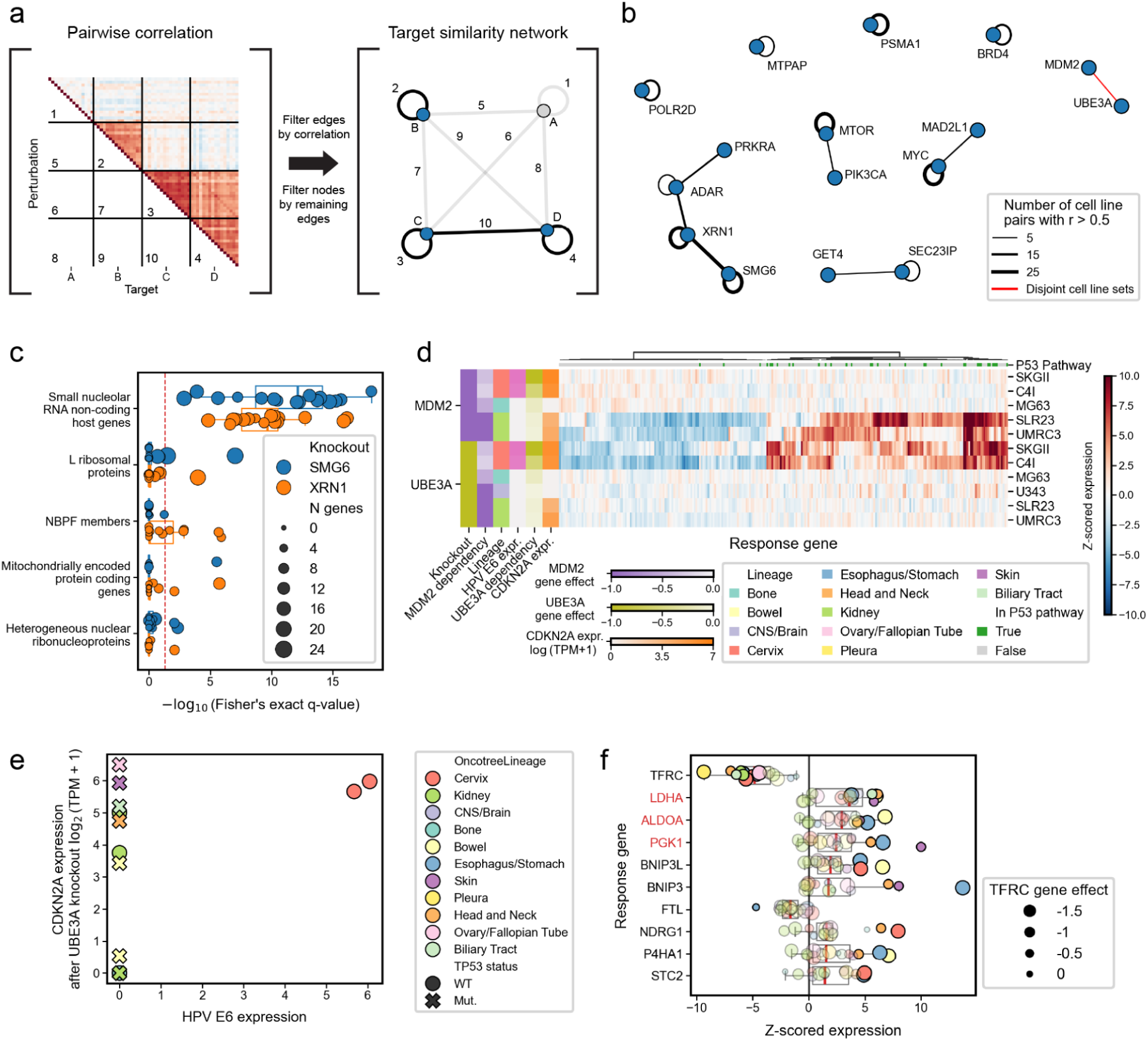
Context-independent and context-dependent gene relationships across an integrated Perturb-seq dataset. **a.** Workflow to produce a context-aware gene similarity graph from pairwise correlations in transcriptional response. **b.** Similarity graph after filtering edges to pairs of knockouts with at least 4 pairs of cell lines inducing responses with high correlation (r > 0.5). **c.** Gene set enrichment across HGNC gene families within the 100 most strongly dysregulated genes in *SMG6*- and *XRN1*-knockout cell lines. **d.** Heatmap of z-scores in response to *MDM2* or *UBE3A* knockout for genes with significantly differential expression in at least 2 perturbations. **e.** Pseudobulked *CDKN2A* expression and viral E6 expression in *TP53*-mutant cell lines with *UBE3A* knockout. **f.** Change in expression for the 10 most strongly differential genes across cell lines following *TFRC* knockout. Genes involved in the glycolysis pathway are labeled in red.

Screening across many contexts allows us to surface relationships that are invisible in any one context. For example, we observed a strong common upregulation of p53 pathway genes following *MDM2* and *UBE3A* knockout, but each knockout caused this phenotype in a different subset of *TP53* wild-type lines (Figure 5d, Supplementary Figure 6c). Both genes encode negative regulators of the tumor suppressor protein p53, with *MDM2* encoding the canonical repressor which ubiquitinates p53 and *UBE3A* encoding another ubiquitin ligase that interacts with the E6 protein found in human papilloma virus (HPV) to bind p53 [38,39]. The models which responded to *UBE3A* knockout (number of differentially expressed genes > 0) are HPV-associated cervical cell lines SKGII and C4I expressing viral E6 and *CDKN2A*, the gene which encodes the MDM2-interfering protein p14ARF (Figure 5e) [40]. *CDKN2A* expression may explain why MDM2 does not compensate for loss of *UBE3A* in SKGII and C4I. Conversely, *UBE3A* cannot compensate for loss of *MDM2* in UMRC3 and SLR23, which lack the viral E6 protein.

Transcriptome-wide correlations can only be large if many genes are dysregulated after a knockout. Some genes have functions which do not induce a broad enough transcriptional phenotype to drive correlations in our network, but nonetheless exhibit consistent and specific effects that illuminate their biology. For example, the top four genes following *TFRC* knockout included downregulation of *TFRC* itself and upregulation of three enzymes in the glycolysis pathway, especially in sensitive lines (Figure 5f, Hallmark GLYCOLYSIS *q* = 6.3 x 10^−7^). *TFRC* and *TFR2* encode the two receptors which are responsible for importing transferrin-bound iron, a critical pathway for iron uptake in cells. Though *TFRC* itself has no direct role in glycolysis, Kumar et al. found that GAPDH moonlights as both a glycolytic enzyme and a transferrin receptor, and that GAPDH membrane localization increases with reduced *TFRC* and *TFR2* expression [41,42]. We observed moderate correlation between *TFRC* dependency in DepMap and change in *GAPDH* expression after TFRC knockout in Perturb-seq (Pearson’s *r* = −0.47, *p* = 0.07), Supplementary Figure 6d). The reallocation of limited GAPDH protein from glycolysis to membrane transferrin function, necessary to compensate for impaired iron import, may force the cell to compensate for GAPDH’s lost glycolytic function in turn by increasing expression of other glycolytic enzymes.

### *IER3IP1* knockout induces oxidative stress in dependent cell lines

To identify less extreme but consistently similar responses between target genes, we computed an alternative network of gene similarity requiring a lower threshold of correlation but spanning more cell lines (Pearson’s r > 0.35, cell line pairs ≥ 10, Figure 6a). This revealed another cluster of genes centralized around *SEC23IP*, whose functions converge around the endoplasmic reticulum (ER) and the Golgi apparatus. *SCYL1*, *SEC23IP*, *IER3IP1*, *SLC39A9*, and *GET4* knockouts induced correlated transcriptional responses in a largely overlapping subset of cell lines (Supplementary Figure 7a). These genes show strongly selective essentiality across DepMap, making them potentially attractive therapeutic targets to explore further (Supplementary Figure 7b).

**Figure 6.**
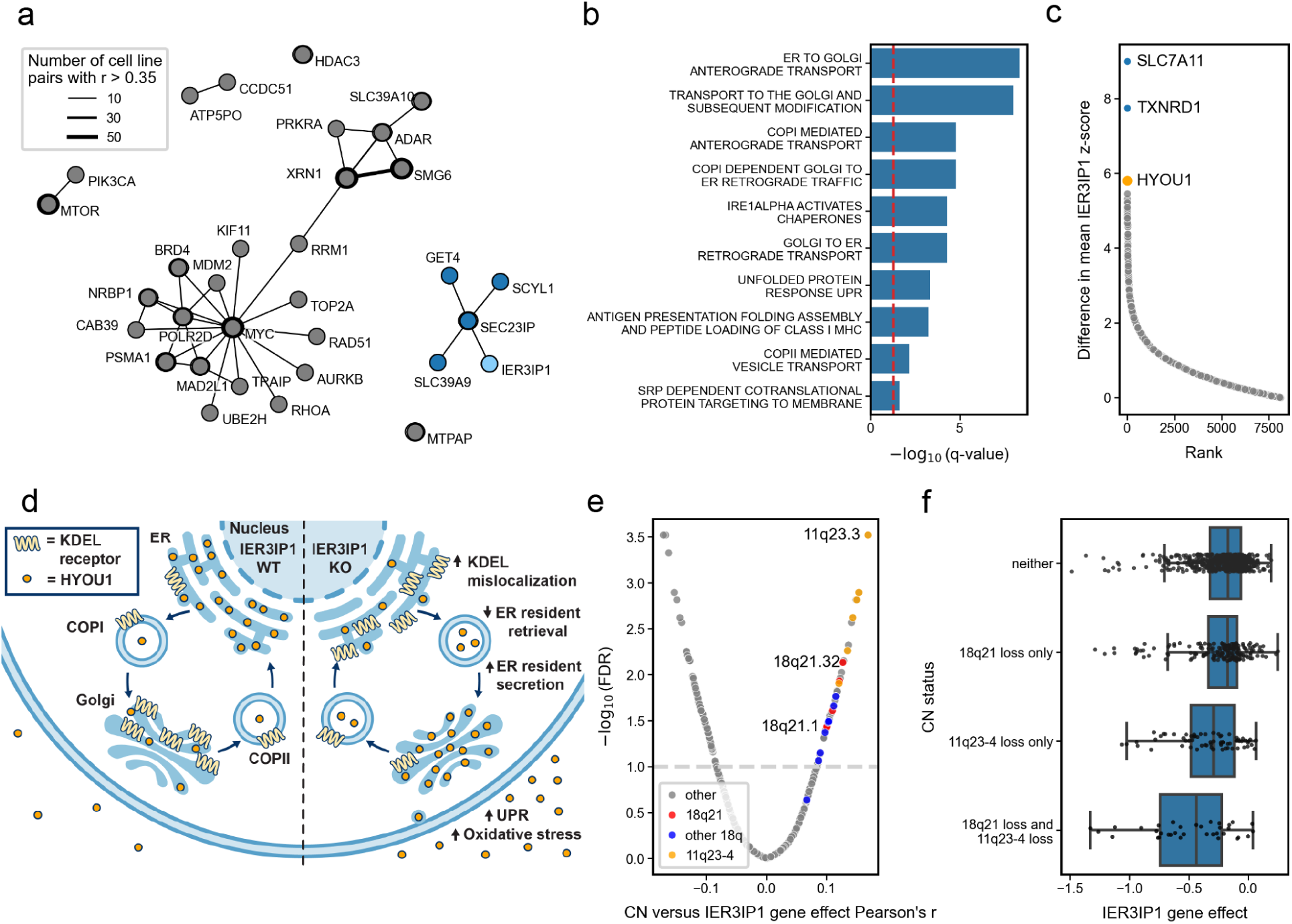
Investigation of ER-golgi transport related KOs. **a.** Similarity graph created by connecting pairs of knockouts with at least 10 spanning pairs of cell lines inducing responses with correlation above r = 0.35. *SEC23IP* cluster containing *SCYL1*, *SEC23IP*, *IER3IP1*, *SLC39A9*, and *GET4* highlighted in blue **b.** Gene set enrichment for shared response genes following KO of *SEC23IP* cluster genes. **c.** Waterfall plot for the difference in mean z-score following *IER3IP1* knockout versus other *SEC23IP* cluster knockouts in dependent cell lines. **d.** Mechanistic model for *IER3IP1* KO resulting in increased *HYOU1* mislocalization. **e**. Volcano plot for Pearson correlation of cytoband copy number with IER3IP1 dependence. **f.** Boxplot showing the relationship between *IER3IP1* dependence and hemizygous loss of cytoband 18q21 and 11q23-4.

We investigated the common response among these knockouts and found consistent upregulation of ER-Golgi transport and IRE1α mediated unfolded protein response (Figure 6b). This included members of the ER to Golgi anterograde and retrograde pathways, including COPI (*COPA*, *COPB2*) and COPII (*SEC31A*, *SEC24A*, *SEC24B*) vesicle coat proteins, as well as several proteins known to circulate between these organelles (*KDELR2*, *KDELR3*, *YIF1A*, *YIPF5*) (Supplementary Fig 7c). The shared response also included *C11orf24*, a gene encoding a poorly characterized protein thought to cycle between the Golgi and plasma membrane via RAB6 positive vesicles [43]. In general, more dependent KO-cell line pairs showed greater upregulation of this shared signature (Supplementary Figure 7d).

In addition to this shared transport and UPR signal, we noted *IER3IP1* dependent cell lines showed stronger upregulation of *SLC7A11, TXNRD1,* and *HYOU1* than other knockouts in the *SEC23IP* cluster (Figure 6c, Supplementary Figure 7e). These three genes are components of *NFE2L2*/NRF2 driven oxidative stress response[44,45]. This hypoxic signature appears to be a direct consequence of *IER3IP1* knockout, as *IER3IP1*-dependent lines did not exhibit higher basal levels of oxidative stress (Supplementary Figure 7f-g). Recently, Anitei et al. demonstrated that loss of *IER3IP1* leads to mislocalization of *LMAN1*/ERGIC53 and *KDELR2* from the Golgi, in turn leading to excessive secretion of ER resident proteins including *HYOU1* (Figure 6d) [46]. In *IER3IP1*-sensitive cells, loss of *HYOU1* likely drives toxic hypoxia.

In order to identify characteristics of IER3IP1 sensitive cell lines, we examined the association of dependence to baseline genomic features. The connection to *HYOU1* illuminates a pattern in the larger Achilles dataset: *IER3IP1* dependence is partially explained by hemizygous deletion of cytobands 18q21.1-32 and 11q23-4 (Mann-Whitney *p* = 3.28 x 10^−6^, Figure 6e,f). These genomic regions contain *IER3IP1* itself (18q21.1), *LMAN1* (18q21.32), and *HYOU1* (11q23.3) – suggesting a potential synthetic lethality between co-deletion of these genes and further *IER3IP1* suppression. This points to a therapeutic opportunity, as loss of 18q is a common event in cancer, and in particular, the loss of the 18q21 region has been repeatedly associated with tumor progression and poor prognosis [47–49]. While additional experimental work and biomarker refinement would be necessary to validate *IER3IP1*’s potential as a drug target, this example demonstrates the mechanistic insights Perturb-seq can offer.

## DISCUSSION

Perturb-seq has been established as an efficient and scalable approach for understanding gene function, linking transcriptional phenotypes to precise genetic knockouts simultaneously for hundreds to thousands of genes [7,10,11,50,51]. However, our ability to interrogate gene function in only one cell line may be limited by the variation in expression and gene essentiality across different models. We have generated an integrated Perturb-seq dataset across 16 different cancer cell lines in 11 lineages, which we have used to find new insights into potential targets. We developed additional quality control methods to remove Cas9-induced genetic artifacts and verified that we recovered expected knockout responses at the level of individual genes.

In cancer biology, Perturb-seq is an exciting means to investigate target biology at scale. Among the targets in our library, we found common markers of cell stress and decomposed cell viability loss into specific phenotypes. We constructed functional networks of genes with similar response phenotypes. Finally, we illustrated the power of Perturb-seq for hypothesis generation. For example, the transferrin receptor *TFRC* causes upregulation of a glycolysis module, especially in sensitive lines, suggesting that iron intake and glycolysis compete for the moonlighting protein *GAPDH* in these lines; *GAPDH* expression is protective against *TFRC* dependency in DepMap. Multiple knockouts upregulated anterograde transport genes. Of these, *IER3IP1* stood out for also strongly upregulating hypoxia response genes in sensitive lines, including *HYOU1*; previous literature showed that *IER3IP1* loss causes secretion of *HYOU1* from the endoplasmic reticulum, and hemizygous loss of both the cytobands containing *IER3IP1* and *HYOU1* sensitize cells to *IER3IP1* knockout in DepMap. For both *TFRC* and *IER3IP1*, Perturb-seq excavated and illuminated overlooked relationships in the Dependency Map data.

A major challenge in this work was the limited number of target genes covered. Lacking a large library of knockouts with well-understood effects, we had to interpret each phenotype manually. Genome-scale assays across more cell contexts will help to scale the interpretation of target response. Additionally, we found that the number of recovered UMIs is the critical metric for observable signal in the dataset, and suggest others in the field to focus on this measure and not only total cells screened. As sequencing costs have rapidly fallen, we will aim for at least 5 million UMIs/knockout in future experiments.

## CONCLUSIONS

In this study, we demonstrate the feasibility of large-scale Cas9 Perturb-seq screening. We establish robust experimental and quality control methods that generalize across a diverse collection of cancer cell lines. In spite of the limited library of 100 targets, we observe expected and novel post-perturbational phenotypes. By combining expression phenotypes with the pan-cancer DepMap corpus, we gained new insights into the mechanism of several compelling dependencies. Future genome-scale Perturb-seq in diverse cancer contexts would enrich understanding of both oncology and general cell biology.

## METHODS

### Cancer cell lines and Cas9 engineering

We selected cell lines previously profiled by DepMap [2] with a CRISPR viability screen, whole exome or genome sequencing, and RNA sequencing. Parental cell lines used were sourced from DepMap Project and Cancer Cell Line Encyclopedia (CCLE) banks at the Broad Institute. The original acquisition of all cancer models was through certified repositories or academic collaborators. Specific details regarding each line’s origin are listed in Supplementary Table 2 and can also be accessed via the DepMap Portal (https://depmap.org/portal/). All cancer cell lines were cultured in RPMI-1640 media containing 10% FBS and 100 units/ml penicillin-streptomycin and engineered to constitutively express Cas9 through lentiviral transduction of pLX311-Cas9 vector (Addgene #96924). Routine validation, including Cas9 activity assay, short tandem repeats (STR) profiling and mycoplasm contamination testing (ABM, G238), as conducted across all lines to ensure quality and authenticity.

Cas9-expressing cells were maintained under blasticidin selection, initially at 10 μg/mL for one week followed by 5 μg/mL for maintenance. Cas9 activity was evaluated by comparing viability following transduction with an sgRNA targeting the pan-essential gene *SF3B1* to a negative-control sgRNA (Chr2-2) across multiple viral doses. Relative viability was assessed 5–6 days after transduction.

Puromycin sensitivity was determined independently for each cell line by testing concentrations ranging from 0–8 μg/mL and selecting the lowest concentration that resulted in near-complete elimination of untransduced cells following 72 hours of treatment.

The appropriate viral dose for the Perturb-seq library was determined empirically for each cell line. Cells were spinfected with a range of library virus volumes at 900 × g for 1.5 h at 37°C in parallel cultures with and without puromycin selection. Three days after initiation of selection, viable cell numbers were compared between selected and unselected cultures. The viral dose producing approximately 15% relative viability following puromycin selection was selected for the screen, corresponding to the desired low-MOI condition. Cells transduced at the selected dose were replated following selection and harvested on day 6 for single-cell capture.

### Perturb-seq screening

To keep the 100 gene library compact without compromising the efficiency of the CRISPR knockout, we designed a library of dual guide constructs with two constructs per gene (4 guides per gene total). Lentiviral constructs were designed to contain puromycin under control of EF1a, one guide under control of a U6 promoter, and a second guide targeting the same gene under control of an H1 promoter. For the 10 gene library for the deep rescreens of KMRC20 and UMRC3, we used the CROP-seq-opti vector (Addgene #106280) with 4 single guide constructs per gene. In both libraries, Cas9 guides for our targets were designed using Rule Set 2 [52]. Cells were infected with several titers to achieve an MOI of 0.2, then selected under puromycin for 96 hours. We grew cells for 6 days after infection to allow for guide transduction and activation of Cas9 cutting activity.

### Single-cell RNA sequencing and processing

Generation of single-cell samples and subsequent library preparation were performed according to the 10x Genomics Single Cell 3’ v3.1 Dual Index Gene Expression with Feature Barcode protocol. After sequencing the samples using the Illumina NovaSeq system, reads were mapped and quantified using CellRanger version 6.0.1, with the *–scaffold_sequence*:

### GTTTAAGAGCTATG

Gene expression reads were aligned to hg38 using the STAR algorithm, as performed by CellRanger, then quantified according to the number of unique molecular identifiers (UMIs) associated with the reads mapping to each gene. CRISPR reads were quantified by fitting a mixture model between guide-expressing cells and cells with ambient sgRNA reads, reporting UMI counts for only cells likely to be guide-expressing. This method is also performed as part of the CellRanger pipeline [53].

Deep rescreens of KMRC20 and UMRC3 were performed with 10x Genomics GEM-X Flex v1 Gene Expression, with custom probes for guide detection. Data was processed on 10x Genomics Cloud using Cell Ranger Multi v9.0.1 with Chromium Human Transcriptome Probe Set v1.1.0 and a custom Feature Reference for CRISPR Guide Capture, producing both gene expression and guide level UMI counts.

### SCEPTRE configuration

To test for differential expression, we applied SCEPTRE in its low-MOI setting. SCEPTRE is designed to perform analysis of Perturb-seq data from quality control through differential expression analysis, using permutation testing to assess differential gene expression in response to CRISPR knockout [24,54]. We used its “maximum” method for assignment of guides, requiring a minimum of 5 reads in the maximal guide and requiring that the maximum guide comprises 80% of the sgRNA reads in a cell. For cell-wise quality control, we kept cells within the 1st and 99th quantiles of both the number of genes detected and the total number of UMIs in a cell. We excluded cells with mitochondrial gene expression comprising more than 25% of the total reads. For knockout-response pair quality control, we required at least 7 nonzero counts of each gene in the knockout and control conditions. We ran differential analysis in verbose mode to retrieve the test scores used to compute significance values. For each target gene and response gene pair, SCEPTRE produces estimates of log-fold change (LFC), a significance value, and an observed Z-score under its negative binomial model which is compared against a set of random scores to obtain the significance value. Downstream analysis involving differential expression uses Z-scores as effect sizes. Using the reported significance values, we identified differentially expressed genes as having FDR < 0.05 after applying the Benjamini-Hochberg multiple hypothesis correction per perturbation [55].

### Quality control of expression profiles

To measure the fidelity of basal transcriptional profiles in our screens, we pseudobulked negative control (olfactory receptor knockout) cells within each cell line and correlated them against all bulk RNA-seq profiles generated by Depmap, considering only the most highly variable genes’ (*n* = 500) expression [56]. For each cell lines’ pseudobulk the top correlate was the expected bulk RNA-seq profile (0.87 < Pearson’s *r* in matched cell lines < 0.95; −0.19 < Pearson’s *r* in mismatched cell lines < 0.88, Supplementary Figure 3a), indicating that defining expression markers of each cell line were preserved after negative control perturbations.

### Arm loss identification

We predicted arm truncation in individual cells by comparing read coverage along the targeted arm between perturbed cells and control cells. For each target *t*, we identified all genes located closer to the telomere on the targeted arm using genomic coordinates from hg38. Then, for all cells, we calculated the fraction of its total reads mapping to distal genes, *F_t_*. Using the distribution of *F_t_* values for all cells receiving a control perturbation which target a gene on a different arm than *t*, we selected the 2.5th quantile to use as a lower threshold *T_t_* for expected read coverage at that arm. Individual cells with *F_t_* < *T_t_* were marked as having arm truncation and excluded from all differential expression analyses. Equivalently, we used the 97.5th quantile to identify arm gain.

Pseudobulk expression of genes in cells with arm loss, arm gain, or normal Cas-induced mutation was converted to TPM by summing expression over cells, dividing by the total readcounts, and multiplying by 10^6^. Z-scored expression in individual cells was calculated using the mean and variance over all cells assigned as unperturbed (total CRISPR reads ≤ 1) or cells receiving any control guide.

### High variability single cell analysis

To assess highly variable target genes, we computed for each gene *g* and cell line *c*, the index of dispersion in readcounts across cells receiving any negative control perturbation for each cell line *c*, defined as

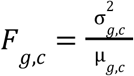

Where *σ^2^_g,c_* represents the variance and µ*_g_*_,*c*_ represents the mean of counts over individual cells. The genes were assessed by their median indices of dispersion *F_g_*_,*c*_ across cell lines. We excluded genes that were detected in fewer than 6 cell lines and genes in the upper 3rd quantile of mean expression.

### Gene set overrepresentation

Gene set overrepresentation analyses were all performed using a Fisher’s exact test with the alternative hypothesis that more genes in the respective gene set are found in the query list of genes than expected by the size of the gene set. For high variance genes, the top 100 genes with the strongest correlation with *S100A8, S100A9, FN1, IFIT2,* and *ISG15* expression across control perturbation cells were chosen as the query list. For genes suspected as having been profiled early and for understanding broad patterns of transcriptional responses to perturbations, we assessed all significantly differentially expressed genes in each perturbation individually.

Gene sets were acquired from the Hallmark (H), Reactome (C2), and Gene Ontology (C5) collections from the Molecular Signatures Database (MSigDB) [33,57]. Gene groups were acquired from the HUGO Gene Nomenclature Committee [58]. Multiple hypothesis correction was performed using the Benjamini-Hochberg procedure [55].

### Correlation network calculation

To broadly assess context-dependent and −independent relationships across genetic knockouts, we constructed a similarity network *G_R,P_*between gene targets based on pairwise Pearson correlation coefficients between transcriptional response profiles for all knockouts in all cell lines (n = 1,322). The correlation analysis yielded at most 120 unique non-trivial correlations between any two genetic targets, spanning all pairs of 16 cell lines. For every pair of target genes, we counted the number of cell line pairs with a correlation coefficient greater than *R*. We used these pair counts to define edges between nodes representing genes, including edges only where there exist at least *P* pairs of cell lines with correlation greater than R. Edges were flagged as representative of context-dependent relationships if two criteria were satisfied:

1. The set of high-correlation pairs exclusively contain pairs spanning different cell lines.
2. The distribution of non-trivial correlations is bimodal. Namely, the distribution is bimodal if the bimodality coefficient is greater than the critical value 5/9, derived from the uniform distribution [59]

We created two networks, *G_0.5,4_* and *G_0.35,10_*, designed to capture strong and highly specific similarity and weaker but broad similarity, respectively.

## DECLARATIONS

### Ethics approval and consent to participate

Not applicable.

### Consent for publication

Not applicable.

### Availability of data and materials

The datasets supporting the conclusions of this article, including single-cell expression data, sgRNA readcounts, and downstream analysis, can be accessed on Figshare [60].The code to produce the results and figures in this study is available on github at https://github.com/broadinstitute/perturb-seq-depmap-public with the version used in this article archived on Figshare [61]. For all cell lines, basal expression and genomic profiles were obtained from the DepMap 25Q2 release at depmap.org.

### Authors contributions

FV conceived this study; TS and PB designed the libraries; BP, AS, TS, PB, JR, HL, SW, AA, OO, and KW conducted the experiments; SM, IAB, LW, and WNC performed computational analysis for this project; SM and LW generated the figures; SM, JMD, and IAB wrote the manuscript; JMD, FV and CDC oversaw this project; All authors revised and approved the manuscript.

## Acknowledgments

Not applicable.

## REFERENCES

1. McDonald ER 3rd, de Weck A, Schlabach MR, Billy E, Mavrakis KJ, Hoffman GR, et al. Project DRIVE: A Compendium of Cancer Dependencies and Synthetic Lethal Relationships Uncovered by Large-Scale, Deep RNAi Screening. Cell. 2017;170:577–592.e10.

2. Tsherniak A, Vazquez F, Montgomery PG, Weir BA, Kryukov G, Cowley GS, et al. Defining a Cancer Dependency Map. Cell. 2017;170:564–576.e16.

3. Behan FM, Iorio F, Picco G, Gonçalves E, Beaver CM, Migliardi G, et al. Prioritization of cancer therapeutic targets using CRISPR-Cas9 screens. Nature. 2019;568:511–6.

4. Pacini C, Duncan E, Gonçalves E, Gilbert J, Bhosle S, Horswell S, et al. A comprehensive clinically informed map of dependencies in cancer cells and framework for target prioritization. Cancer Cell. 2024;42:301–316.e9.

5. Arafeh R, Shibue T, Dempster JM, Hahn WC, Vazquez F. The present and future of the Cancer Dependency Map. Nat Rev Cancer. 2025;25:59–73.

6. Datlinger P, Rendeiro AF, Boenke T, Senekowitsch M, Krausgruber T, Barreca D, et al. Ultra-high-throughput single-cell RNA sequencing and perturbation screening with combinatorial fluidic indexing. Nat Methods. 2021;18:635–42.

7. Datlinger P, Rendeiro AF, Schmidl C, Krausgruber T, Traxler P, Klughammer J, et al. Pooled CRISPR screening with single-cell transcriptome readout. Nat Methods. 2017;14:297–301.

8. Mimitou EP, Cheng A, Montalbano A, Hao S, Stoeckius M, Legut M, et al. Multiplexed detection of proteins, transcriptomes, clonotypes and CRISPR perturbations in single cells. Nat Methods. 2019;16:409–12.

9. Wessels H-H, Méndez-Mancilla A, Hao Y, Papalexi E, Mauck WM 3rd, Lu L, et al. Efficient combinatorial targeting of RNA transcripts in single cells with Cas13 RNA Perturb-seq. Nat Methods. 2023;20:86–94.

10. Replogle JM, Saunders RA, Pogson AN, Hussmann JA, Lenail A, Guna A, et al. Mapping information-rich genotype-phenotype landscapes with genome-scale Perturb-seq. Cell. 2022;185:2559–2575.e28.

11. Adamson B, Norman TM, Jost M, Cho MY, Nuñez JK, Chen Y, et al. A Multiplexed Single-Cell CRISPR Screening Platform Enables Systematic Dissection of the Unfolded Protein Response. Cell. 2016;167:1867–1882.e21.

12. Schraivogel D, Gschwind AR, Milbank JH, Leonce DR, Jakob P, Mathur L, et al. Targeted Perturb-seq enables genome-scale genetic screens in single cells. Nat Methods. 2020;17:629–35.

13. Replogle JM, Norman TM, Xu A, Hussmann JA, Chen J, Cogan JZ, et al. Combinatorial single-cell CRISPR screens by direct guide RNA capture and targeted sequencing. Nat Biotechnol. 2020;38:954–61.

14. Norman TM, Horlbeck MA, Replogle JM, Ge AY, Xu A, Jost M, et al. Exploring genetic interaction manifolds constructed from rich single-cell phenotypes. Science. 2019;365:786–93.

15. Santinha AJ, Klingler E, Kuhn M, Farouni R, Lagler S, Kalamakis G, et al. Transcriptional linkage analysis with in vivo AAV-Perturb-seq. Nature. 2023;622:367–75.

16. Jin X, Simmons SK, Guo A, Shetty AS, Ko M, Nguyen L, et al. In vivo Perturb-Seq reveals neuronal and glial abnormalities associated with autism risk genes. Science. 2020;370:eaaz6063.

17. Zheng X, Wu B, Liu Y, Simmons SK, Kim K, Clarke GS, et al. Massively parallel in vivo Perturb-seq reveals cell-type-specific transcriptional networks in cortical development. Cell. 2024;187:3236–3248.e21.

18. Corsello SM, Nagari RT, Spangler RD, Rossen J, Kocak M, Bryan JG, et al. Discovering the anti-cancer potential of non-oncology drugs by systematic viability profiling. Nat Cancer. 2020;1:235–48.

19. Garnett MJ, Edelman EJ, Heidorn SJ, Greenman CD, Dastur A, Lau KW, et al. Systematic identification of genomic markers of drug sensitivity in cancer cells. Nature. 2012;483:570–5.

20. Barras D. BRAF mutation in colorectal cancer: An update. Biomark Cancer. 2015;7:9–12.

21. Xu T, Wang X, Wang Z, Deng T, Qi C, Liu D, et al. Molecular mechanisms underlying the resistance of BRAF V600E-mutant metastatic colorectal cancer to EGFR/BRAF inhibitors. Ther Adv Med Oncol. 2022;14:17588359221105022.

22. Wykosky J, Fenton T, Furnari F, Cavenee WK. Therapeutic targeting of epidermal growth factor receptor in human cancer: successes and limitations. Chin J Cancer. 2011;30:5–12.

23. Krill-Burger JM, Dempster JM, Borah AA, Paolella BR, Root DE, Golub TR, et al. Partial gene suppression improves identification of cancer vulnerabilities when CRISPR-Cas9 knockout is pan-lethal. Genome Biol. 2023;24:192.

24. Barry T, Mason K, Roeder K, Katsevich E. Robust differential expression testing for single-cell CRISPR screens at low multiplicity of infection. Genome Biol. 2024;25:124.

25. Lazar NH, Celik S, Chen L, Fay MM, Irish JC, Jensen J, et al. High-resolution genome-wide mapping of chromosome-arm-scale truncations induced by CRISPR-Cas9 editing. Nat Genet. 2024;56:1482–93.

26. Zuccaro MV, Xu J, Mitchell C, Marin D, Zimmerman R, Rana B, et al. Allele-specific chromosome removal after Cas9 cleavage in human embryos. Cell. 2020;183:1650–1664.e15.

27. Nahmad AD, Reuveni E, Goldschmidt E, Tenne T, Liberman M, Horovitz-Fried M, et al. Frequent aneuploidy in primary human T cells after CRISPR-Cas9 cleavage. Nat Biotechnol. 2022;40:1807–13.

28. Papathanasiou S, Markoulaki S, Blaine LJ, Leibowitz ML, Zhang C-Z, Jaenisch R, et al. Whole chromosome loss and genomic instability in mouse embryos after CRISPR-Cas9 genome editing. Nat Commun. 2021;12:5855.

29. Malnic B, Godfrey PA, Buck LB. The human olfactory receptor gene family. Proc Natl Acad Sci U S A. 2004;101:2584–9.

30. Murakami-Tonami Y, Ikeda H, Yamagishi R, Inayoshi M, Inagaki S, Kishida S, et al. SGO1 is involved in the DNA damage response in MYCN-amplified neuroblastoma cells. Sci Rep. 2016;6:31615.

31. Bhat GR, Sethi I, Sadida HQ, Rah B, Mir R, Algehainy N, et al. Cancer cell plasticity: from cellular, molecular, and genetic mechanisms to tumor heterogeneity and drug resistance. Cancer Metastasis Rev. 2024;43:197–228.

32. Hsiao LL, Dangond F, Yoshida T, Hong R, Jensen RV, Misra J, et al. A compendium of gene expression in normal human tissues. Physiol Genomics. 2001;7:97–104.

33. Liberzon A, Birger C, Thorvaldsdóttir H, Ghandi M, Mesirov JP, Tamayo P. The Molecular Signatures Database (MSigDB) hallmark gene set collection. Cell Syst. 2015;1:417–25.

34. Wang S, Song R, Wang Z, Jing Z, Wang S, Ma J. S100A8/A9 in inflammation. Front Immunol. 2018;9:1298.

35. Ashburner M, Ball CA, Blake JA, Botstein D, Butler H, Cherry JM, et al. Gene ontology: tool for the unification of biology. The Gene Ontology Consortium. Nat Genet. 2000;25:25–9.

36. Gene Ontology Consortium, Aleksander SA, Balhoff J, Carbon S, Cherry JM, Drabkin HJ, et al. The Gene Ontology knowledgebase in 2023. Genetics [Internet]. 2023;224. Available from: 10.1093/genetics/iyad031

37. Lykke-Andersen S, Chen Y, Ardal BR, Lilje B, Waage J, Sandelin A, et al. Human nonsense-mediated RNA decay initiates widely by endonucleolysis and targets snoRNA host genes. Genes Dev. 2014;28:2498–517.

38. Talis AL, Huibregtse JM, Howley PM. The role of E6AP in the regulation of p53 protein levels in human papillomavirus (HPV)-positive and HPV-negative cells. J Biol Chem. 1998;273:6439–45.

39. Sailer C, Offensperger F, Julier A, Kammer K-M, Walker-Gray R, Gold MG, et al. Structural dynamics of the E6AP/UBE3A-E6-p53 enzyme-substrate complex. Nat Commun. 2018;9:4441.

40. Hengstermann A, Linares LK, Ciechanover A, Whitaker NJ, Scheffner M. Complete switch from Mdm2 to human papillomavirus E6-mediated degradation of p53 in cervical cancer cells. Proc Natl Acad Sci U S A. 2001;98:1218–23.

41. Kumar S, Sheokand N, Mhadeshwar MA, Raje CI, Raje M. Characterization of glyceraldehyde-3-phosphate dehydrogenase as a novel transferrin receptor. Int J Biochem Cell Biol. 2012;44:189–99.

42. Sheokand N, Malhotra H, Kumar S, Tillu VA, Chauhan AS, Raje CI, et al. Moonlighting cell-surface GAPDH recruits apotransferrin to effect iron egress from mammalian cells. J Cell Sci. 2014;127:4279–91.

43. Fraisier V, Kasri A, Miserey-Lenkei S, Sibarita J-B, Nair D, Mayeux A, et al. C11ORF24 is a novel type I membrane protein that cycles between the Golgi apparatus and the plasma membrane in Rab6-positive vesicles. PLoS One. 2013;8:e82223.

44. Zhong Z, Zhang C, Ni S, Ma M, Zhang X, Sang W, et al. NFATc1-mediated expression of SLC7A11 drives sensitivity to TXNRD1 inhibitors in osteoclast precursors. Redox Biol. 2023;63:102711.

45. Rao S, Oyang L, Liang J, Yi P, Han Y, Luo X, et al. Biological function of HYOU1 in tumors and other diseases. Onco Targets Ther. 2021;14:1727–35.

46. Anitei M, Bruno F, Valkova C, Dau T, Cirri E, Mestres I, et al. IER3IP1-mutations cause microcephaly by selective inhibition of ER-Golgi transport. Cell Mol Life Sci. 2024;81:334.

47. Cai T, Mondaini N, Tiscione D, Dal Canto M, Santi R, Bartoletti R, et al. Loss of heterozygosis on chromosome 18q21-23 and muscle-invasive bladder cancer natural history. Oncol Lett. 2015;10:2569–73.

48. Jernvall P, Mäkinen MJ, Karttunen TJ, Mäkelä J, Vihko P. Loss of heterozygosity at 18q21 is indicative of recurrence and therefore poor prognosis in a subset of colorectal cancers. Br J Cancer. 1999;79:903–8.

49. Frank CJ, McClatchey KD, Devaney KO, Carey TE. Evidence that loss of chromosome 18q is associated with tumor Progression1. Cancer Res. 1997;57:824–7.

50. Dixit A, Parnas O, Li B, Chen J, Fulco CP, Jerby-Arnon L, et al. Perturb-Seq: Dissecting Molecular Circuits with Scalable Single-Cell RNA Profiling of Pooled Genetic Screens. Cell. 2016;167:1853–1866.e17.

51. Alda-Catalinas C, Bredikhin D, Hernando-Herraez I, Santos F, Kubinyecz O, Eckersley-Maslin MA, et al. A Single-Cell Transcriptomics CRISPR-Activation Screen Identifies Epigenetic Regulators of the Zygotic Genome Activation Program. Cell Syst. 2020;11:25–41.e9.

52. Doench JG, Fusi N, Sullender M, Hegde M, Vaimberg EW, Donovan KF, et al. Optimized sgRNA design to maximize activity and minimize off-target effects of CRISPR-Cas9. Nat Biotechnol. 2016;34:184–91.

53. Zheng GXY, Terry JM, Belgrader P, Ryvkin P, Bent ZW, Wilson R, et al. Massively parallel digital transcriptional profiling of single cells. Nat Commun. 2017;8:14049.

54. Barry T, Wang X, Morris JA, Roeder K, Katsevich E. SCEPTRE improves calibration and sensitivity in single-cell CRISPR screen analysis. Genome Biol. 2021;22:344.

55. Benjamini Y, Hochberg Y. Controlling the false discovery rate: A practical and powerful approach to multiple testing. J R Stat Soc Series B Stat Methodol. 1995;57:289–300.

56. DepMap B. DepMap 24Q4 Public [Internet]. Figshare+; 2024 [cited 2025 Feb 21]. Available from: https://figshare.com/articles/dataset/DepMap_24Q4_Public/27993248/1

57. Subramanian A, Tamayo P, Mootha VK, Mukherjee S, Ebert BL, Gillette MA, et al. Gene set enrichment analysis: a knowledge-based approach for interpreting genome-wide expression profiles. Proc Natl Acad Sci U S A. 2005;102:15545–50.

58. Seal RL, Braschi B, Gray K, Jones TEM, Tweedie S, Haim-Vilmovsky L, et al. Genenames.org: the HGNC resources in 2023. Nucleic Acids Res. 2023;51:D1003–9.

59. Pfister R, Schwarz KA, Janczyk M, Dale R, Freeman JB. Good things peak in pairs: a note on the bimodality coefficient. Front Psychol. 2013;4:700.

60. Ward L. Dissecting context-dependent cancer vulnerabilities using Perturb-seq - Data [Internet]. figshare; 2026 [cited 2026 Aug 24]. Available from: 10.6084/m9.figshare.33273600.v1

61. Ward L. Dissecting context-dependent cancer vulnerabilities using Perturb-seq - Codebase [Internet]. figshare; 2026 [cited 2026 Aug 24]. Available from: 10.6084/m9.figshare.33282171.v1

