## Supplementary Figures for "Dissecting context-dependent cancer vulnerabilities using Perturb-seq"

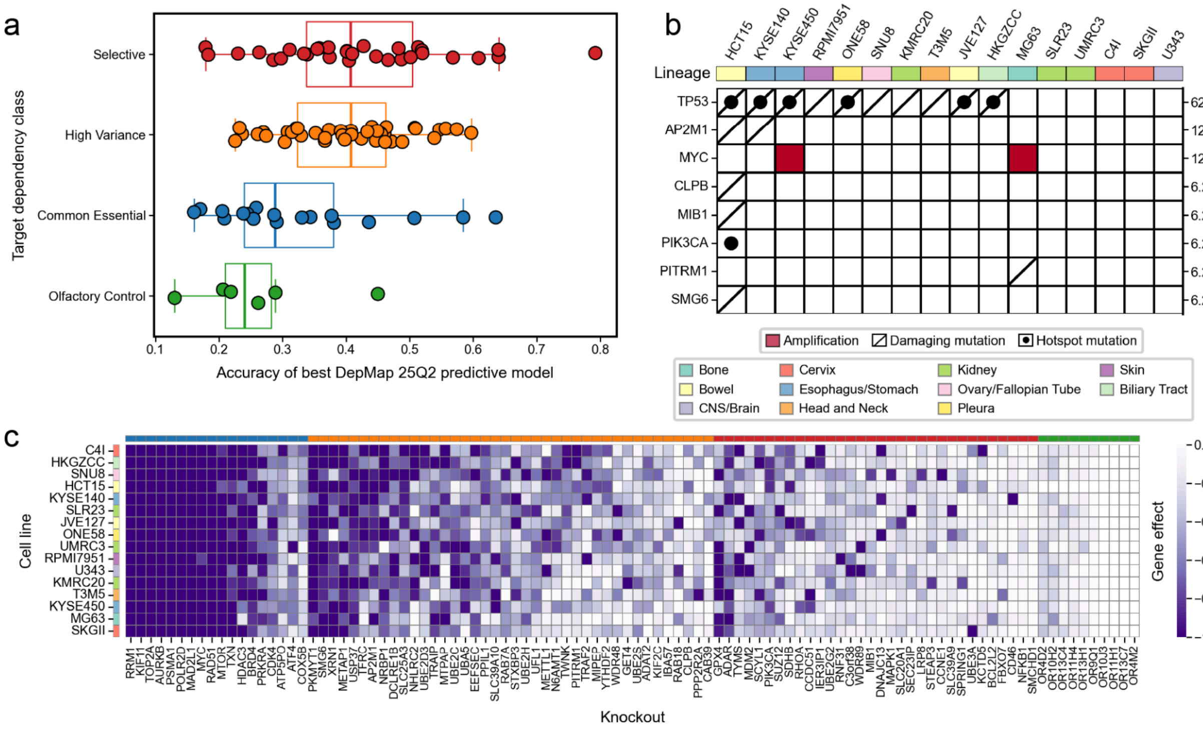


**Supplementary Figure 1.** Characteristics of gene target and cell line panels. **a.** Accuracy (correlation between predicted gene effect and observed gene effect) of the top DepMap predictive model for genes with different categories of dependency profile, based on their dependency distributions across cell lines. Target genes in the guide library are colored by dependency class. **b.** Summary of preexisting genetic alterations in *TP53* and any target included in the sgRNA library. **c.** Heatmap of dependency scores and essentiality patterns across the cell line panel for the genes included in the target library.

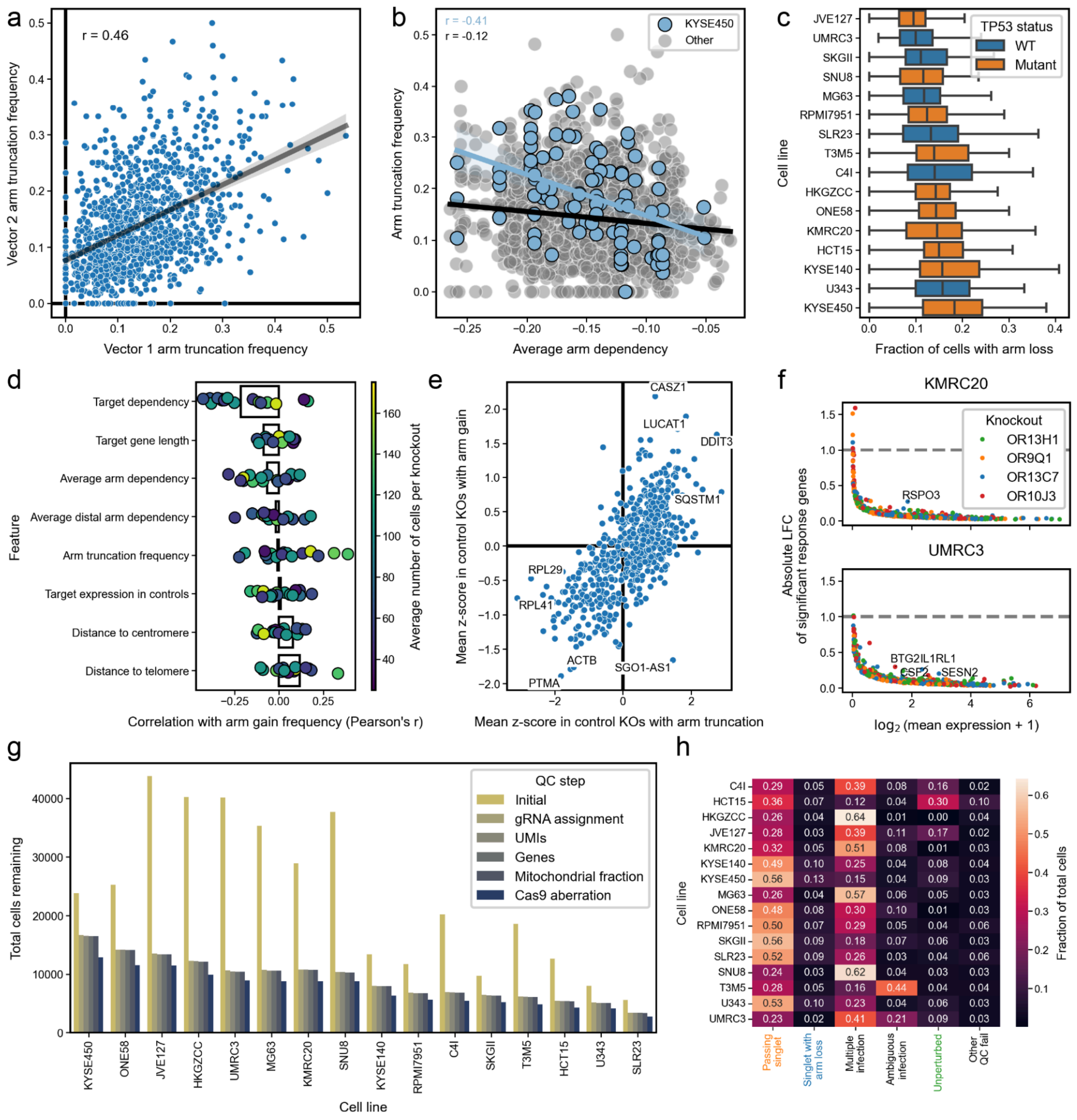


**Supplementary Figure 2.** Characterization of arm truncation events. **a.** Comparison of arm-level truncation frequency caused by different vectors targeting the same gene across the entire library. **b.** Comparison between average dependency of genes on the chromosome arm harboring the target gene and the frequency of arm truncation events detected. Best fit lines for KYSE450 and across all cell lines are overlaid. **c.** Frequency of arm truncation per knockout in different cell lines, colored by *TP53* status. **d.** Correlations between gene-specific properties and frequency of arm gain. Bars represent relationships calculated over all combinations of cell lines and knockouts; points represent relationships calculated over individual cell lines. **e.** Expression differences between control cells with arm truncation and control cells with arm gain relative to normal control cells. **f.** Absolute log-fold change and mean expression of significant responses between olfactory receptor controls and non-cutting controls in deep rescreen of KMRC20 and UMRC3 **g.** Cell recovery per cell line after sequential quality control filters on individual cells. **h.** Summary of cellwise quality control across all Perturb-seq experiments.

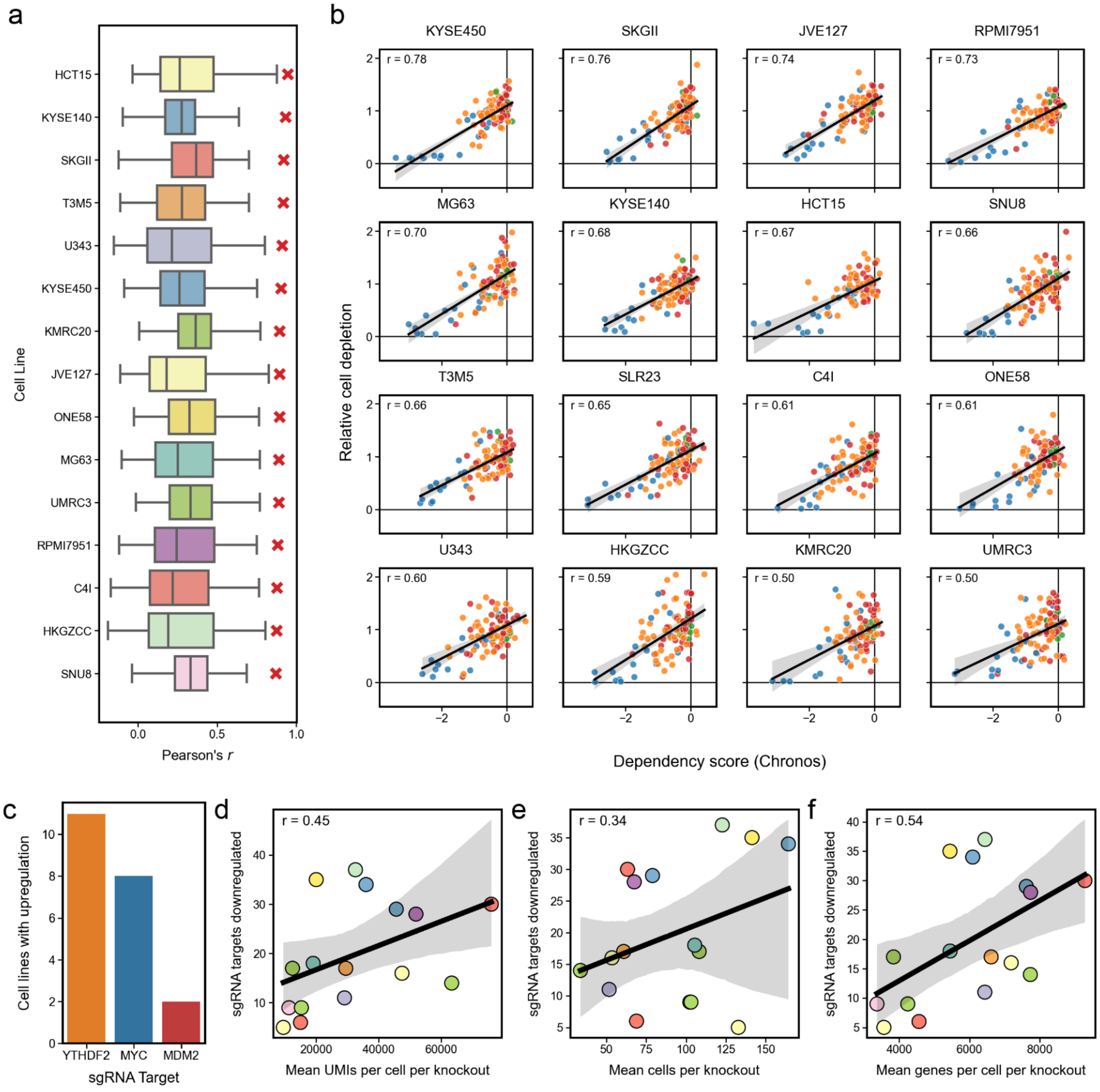


**Supplementary Figure 3.** Additional Perturb-seq quality control metrics. **a.** Correlation between pseudobulk expression in negative control knockouts and bulk expression profiling from DepMap 25Q2. **b.** Cell recovery versus dependency strength for each cell line individually. Colors represent target gene classes. **c.** Gene targets which exhibit significant upregulation after knockout. **d-f.** Sensitivity to detect significant downregulation of the gene target compared to common measures of scRNAseq quality, **d.** average UMIs per cell, **e.** average cells per target, and **f.** average genes detected per cell.

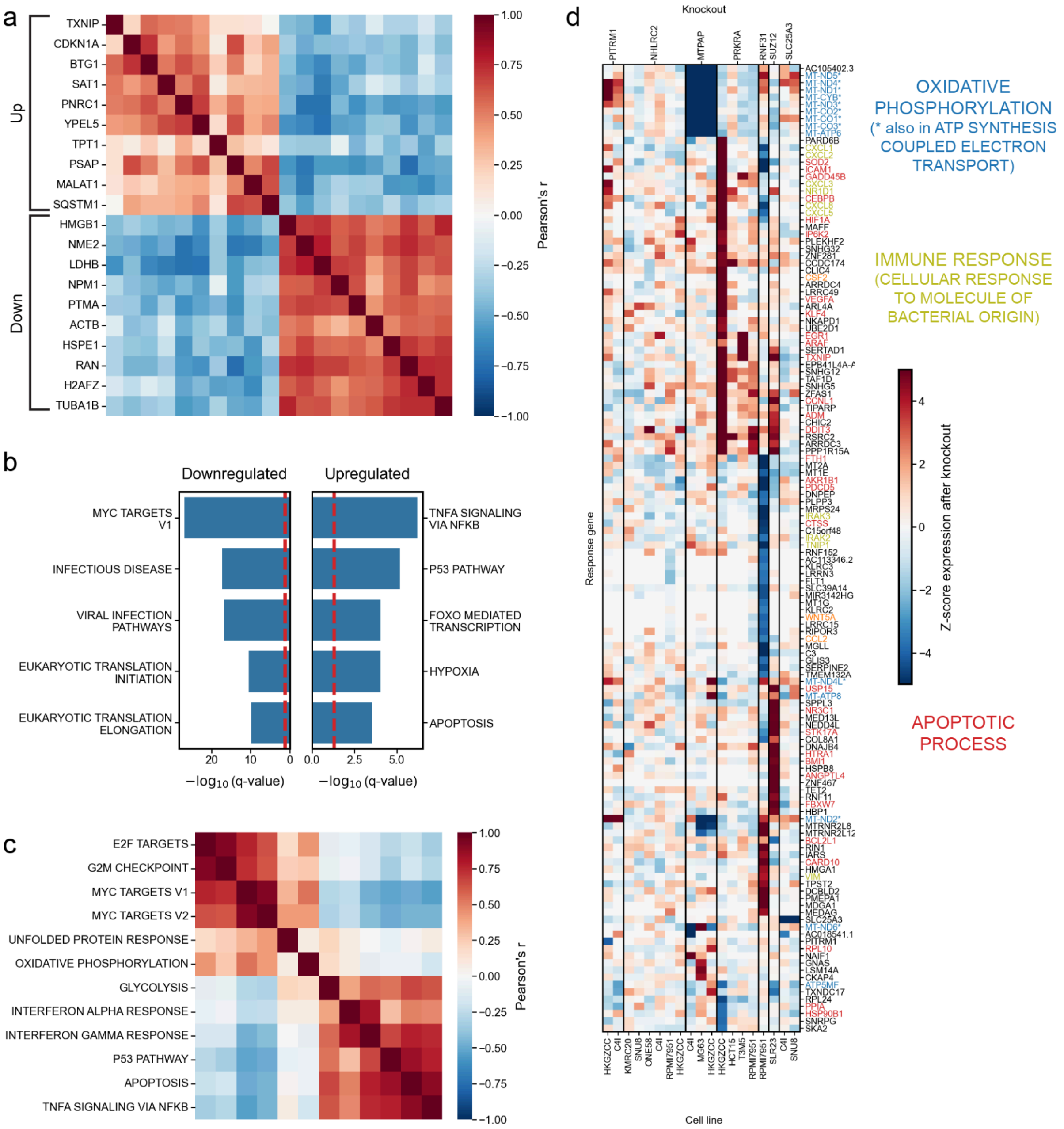


**Supplementary Figure 4.** Common responses to dependent (gene effect < -1) perturbations across 16 cell lines. **a.** Correlation in expression change between the top 10 most commonly upregulated and downregulated genes across dependent knockouts. **b.** Overrepresentation analysis of Hallmark gene sets, Reactome pathways, and HGNC families within the top 50 transcripts most commonly upregulated or downregulated in dependent knockouts. **c.** Correlation between mean change in expression for Hallmark gene sets across all dependent knockouts. **d.** Heatmap of gene expression changes in dependent knockouts lacking downregulation of cell cycle gene expression and exhibiting greater cell recovery than expected by its dependency score. Gene names are colored by membership to Gene Ontology gene sets.

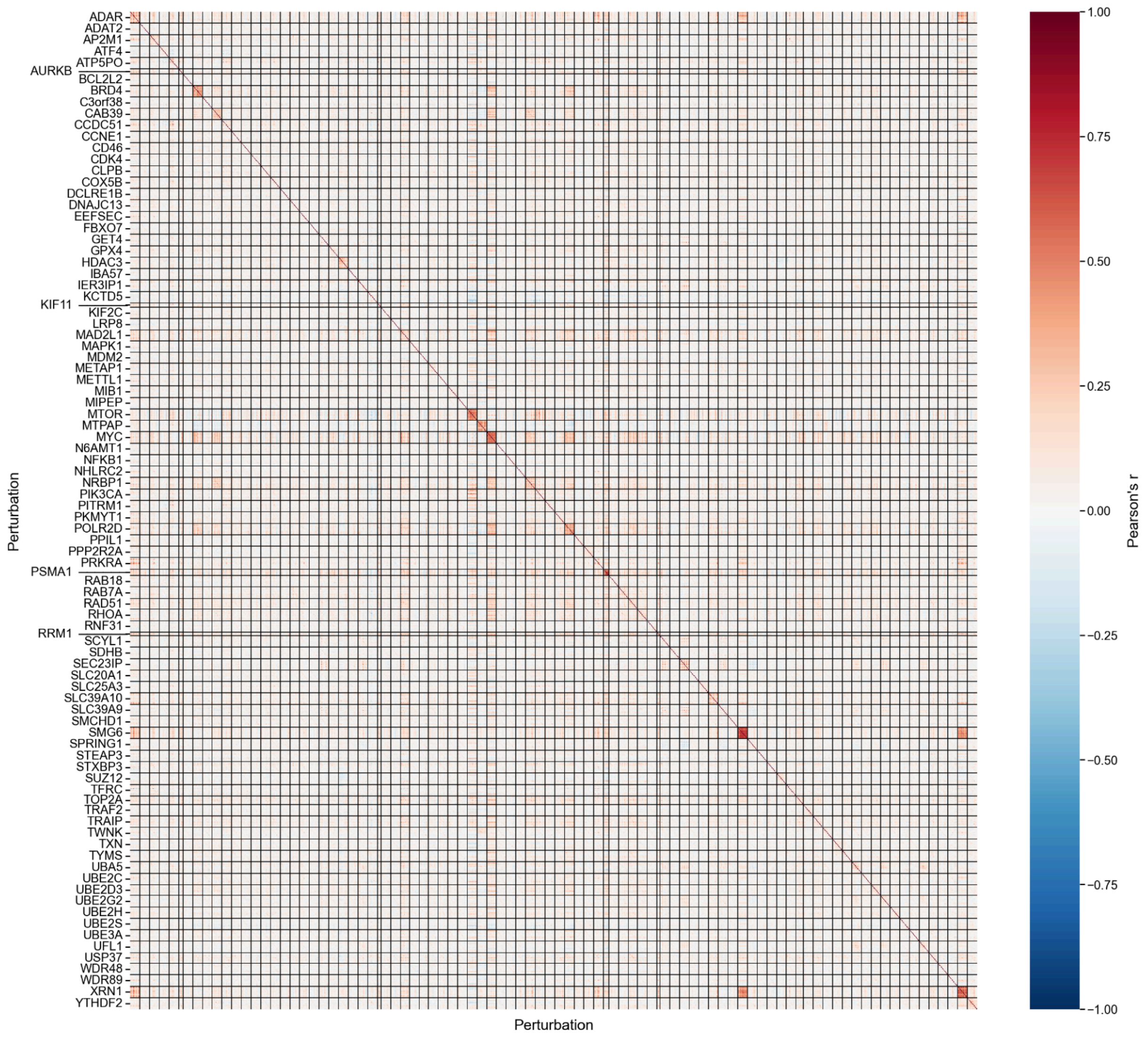


**Supplementary Figure 5.** Correlation between transcriptional changes resulting from all knockouts in all cell lines. Black dividing lines separate gene targets.

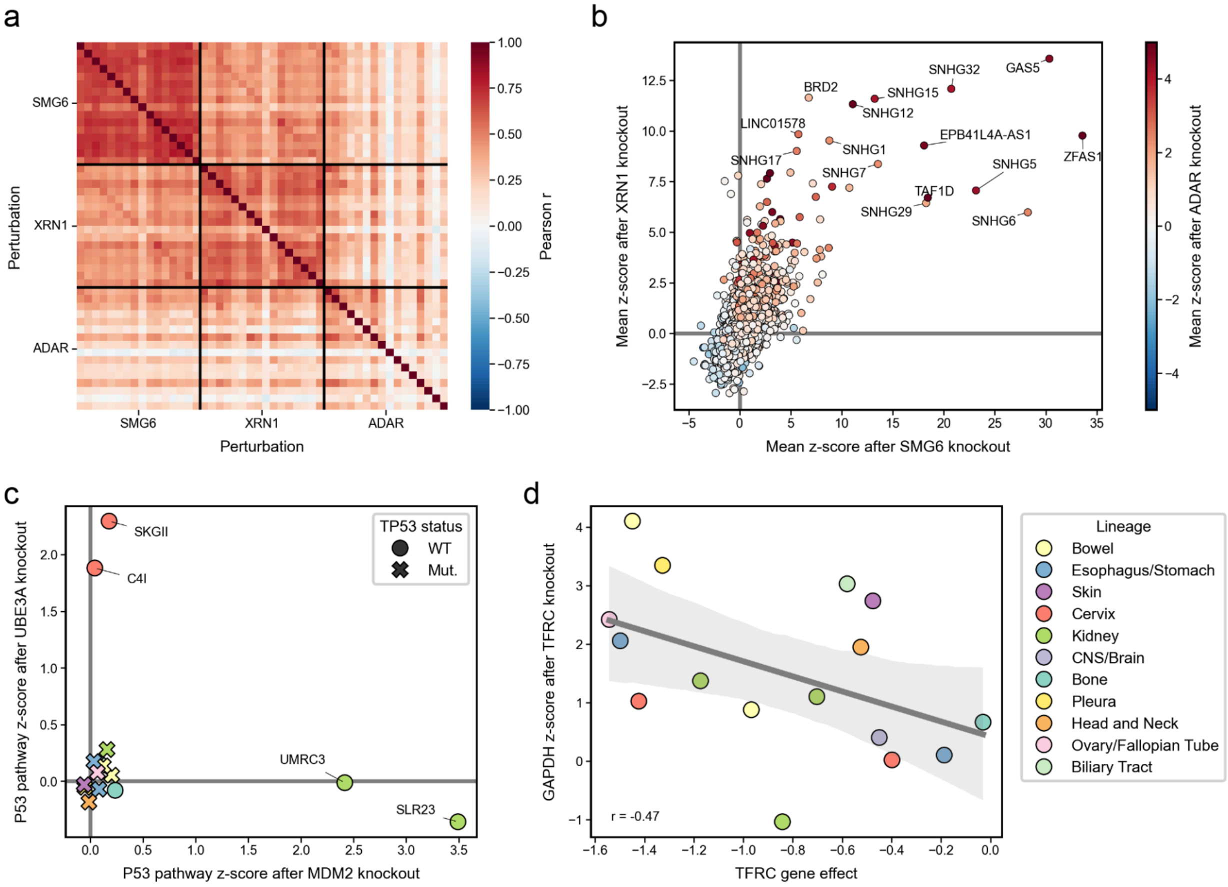


**Supplementary Figure 6.** Expanded vignettes for context-dependent and context-independent responses to gene knockout. **a.** Correlation between all cell lines with *SMG6*, *XRN1*, and *ADAR* knockouts. **b.** Average expression change across all cell lines with *XRN1* and *SMG6* knockouts, colored by average expression change in *ADAR* knockouts. The 10 most extreme genes after *XRN1* and *SMG6* knockout are labeled. **c.** Average change in expression of Hallmark P53_PATHWAY genes per cell line following *MDM2* or *UBE3A* knockout. **d.** Change in expression of *GAPDH* compared to dependency on *TFRC* across all cell lines.

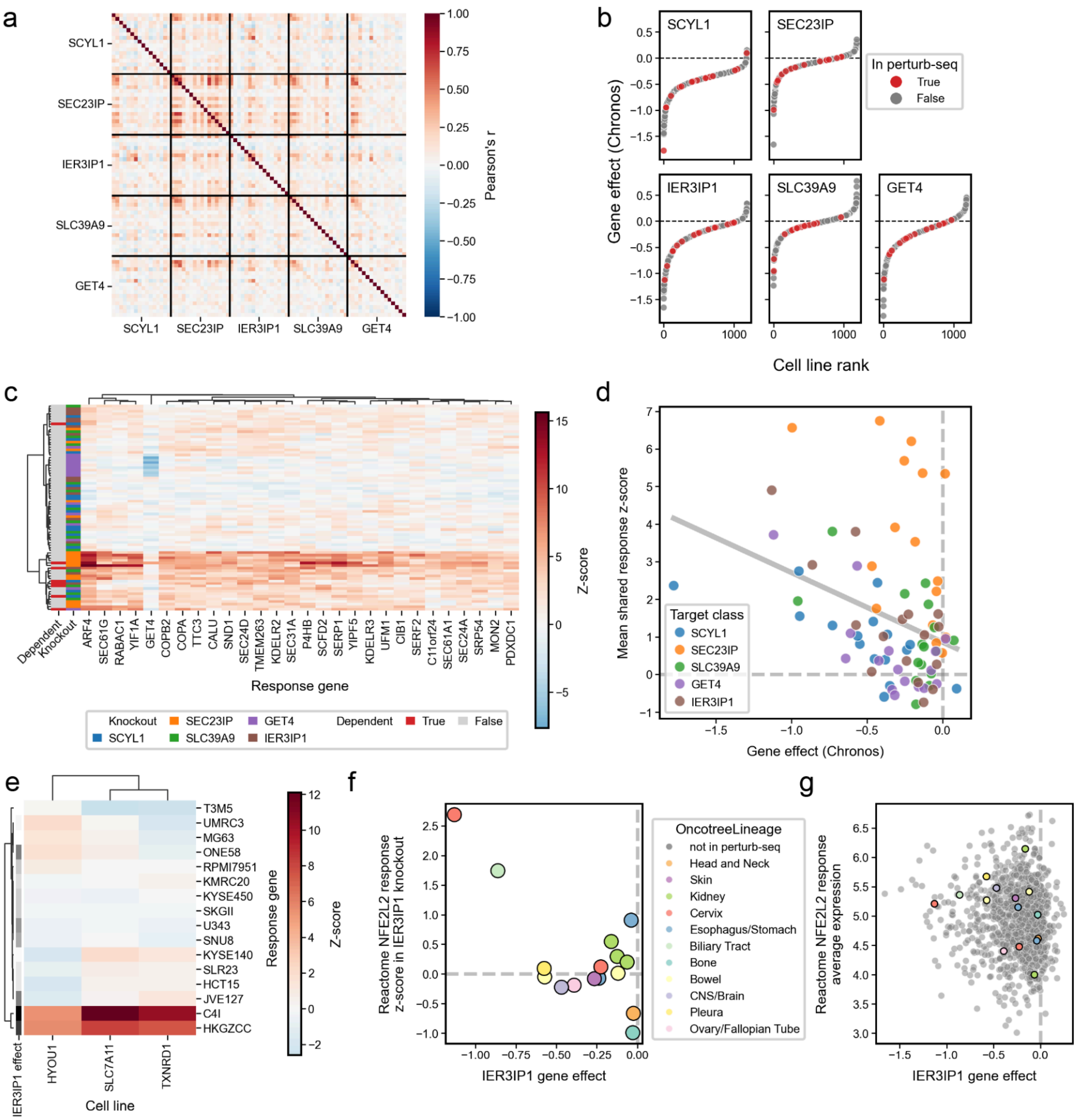


**Supplementary Figure 7.** Further investigation of ER-golgi transport related KOs. **a.** Subset of the correlation heatmap in **6a** for knockouts *SCYL1*, *SEC23IP*, *IER3IP1*, *SLC39A9*, and *GET4*. **b.** Waterfall plots of dependency scores across all DepMap models for genes clustering with *SEC23IP*. Red points indicate cell lines screened with Perturb-seq. **c.** Mean z-score of shared response genes in **6b** versus targets' dependency scores. **d.** Z-scores for the shared response genes in the *SEC23IP* cluster. **e.** Z-scores for *IER3IP1* specific response genes in **6c** for *IER3IP1* KO cells. **f.** Mean z-score of Reactome NFE2L2 response gene set for *IER3IP1* KO cells versus *IER3IP1* dependency. **g.** Average basal bulk RNAseq expression of Reactome NFE2L2 response gene set versus *IER3IP1* dependency.
